# Light/dark and temperature cycling modulate metabolic electron flow in *Pseudomonas aeruginosa* biofilms

**DOI:** 10.1101/2022.02.07.479496

**Authors:** Lisa Juliane Kahl, Kelly N. Eckartt, Diana K. Morales, Alexa Price-Whelan, Lars E. P. Dietrich

## Abstract

Sunlight drives phototrophic metabolism, which affects redox conditions and produces substrates for non-phototrophs. These environmental parameters fluctuate daily due to Earth’s rotation, and non-phototrophic organisms can therefore benefit from the ability to respond to, or even anticipate, such changes. Circadian rhythms, such as daily changes in body temperature, in host organisms can also affect local conditions for colonizing bacteria. Here, we investigated the effects of light/dark and temperature cycling on biofilms of the opportunistic pathogen *Pseudomonas aeruginosa* PA14. We grew biofilms in the presence of a respiratory indicator dye and found that greater dye reduction occurred in biofilm zones that formed during dark intervals and at lower temperatures. This pattern formation occurred with cycling of blue, red, or far-red light, and a screen of mutants representing potential sensory proteins identified two with defects in pattern formation, specifically under red light cycling. We also found that the physiological states of biofilm subzones formed under specific light and temperature conditions were retained during subsequent condition cycling. Light/dark and temperature cycling affected expression of genes involved in primary metabolic pathways and redox homeostasis, including those encoding electron transport chain components. Consistent with this, we found that *cbb*_3_-type oxidases contribute to dye reduction under light/dark cycling conditions. Together, our results indicate that cyclic changes in light exposure and temperature have lasting effects on redox metabolism in biofilms formed by a non-phototrophic, pathogenic bacterium.

**IMPORTANCE:** Organisms that do not obtain energy from light can nevertheless be affected by daily changes in light exposure. Many aspects of animal and fungal physiology fluctuate in response to these changes, including body temperature and the activities of antioxidant and other redox enzymes that play roles in metabolism. Whether redox metabolism is affected by light/dark and temperature cycling in bacteria that colonize such circadian organisms has not been studied in detail. Here we show that growth under light/dark and temperature cycling leads to rhythmic changes in redox metabolism in *Pseudomonas aeruginosa* and identify proteins involved in this response. *P. aeruginosa* is a major cause of healthcare-associated infections and designated as a serious threat by the CDC due to its recalcitrance during treatments. Our findings have the potential to inform therapeutic strategies that incorporate controlled light exposure or consider *P. aeruginosa*’s responses to conditions in the host.

## INTRODUCTION

Sunlight powers most biology and, via its role in phototrophic metabolism, influences cellular and environmental redox potentials. Oxygenic phototrophs in particular, which are thought to have evolved over 2.4 billion years ago (1), use light to produce a strong oxidant (oxygen) that is a major substrate for non-phototrophs (i.e., chemotrophs) and affects the biogeochemical cycling of essential life elements such as sulfur and iron (2, 3). Light and redox therefore have a long-standing relationship in Earth’s history, and because light is only available for a limited portion of each day, redox conditions fluctuate with corresponding regularity (2, 4–6). Though non-phototrophic bacteria do not depend on light as an energy source, they nevertheless can be directly affected, for example, by the impact of light on cellular cofactors, or indirectly affected by the impact of light on the environment. Commensal and pathogenic bacteria can also indirectly experience the effects of light via host circadian rhythms, such as daily fluctuations in body temperature (7–10).

Chemotrophs exhibit responses to visible light exposure as a direct cue (11–15) and also when circadian rhythms, which are controlled by endogenous “clock” mechanisms, are entrained by light cues (16–18). In both cases, studies examining these types of responses have revealed links between redox physiology and light in chemotrophs. In many chemotrophs, substrates donate electrons to central metabolic pathways, which rely on redox reactions to ultimately produce a membrane potential that drives ATP synthesis. Visible light has been shown to affect central metabolism and/or membrane potential in chemotrophic flavobacteria, *Escherichia coli, Bacillus subtilis*, and budding yeast (19–22). For diverse organisms that exhibit light-entrainable circadian physiological rhythms, it has been shown that these rhythms correlate with changes in the redox states of antioxidant/signaling proteins called peroxiredoxins (23). Budding yeast also shows endogenous respiratory oscillations that bear similarities to light-entrainable, endogenous physiological rhythms in other organisms (24), which may hint at an ancestral relationship between redox state and cyclic changes in light exposure. Together, these findings point to fundamental influences of Earth’s fluctuating light conditions on cellular electron-transfer metabolism.

In this study, we examined the physiological response to light/dark and temperature cycling in *Pseudomonas aeruginosa*, a bacterium that causes opportunistic, biofilm-based infections in diverse hosts. *P. aeruginosa* is exposed to light/dark and temperature cycling when it resides close to the surface in soil and aquatic settings and when it colonizes the surfaces of plants and animals, e.g. when it causes leaf or wound infections (25–30). Our results indicate that light/dark and temperature cycling leads to oscillations in routes of metabolic electron flow that are mediated by global changes in transcription. They also suggest that *P. aeruginosa*’s terminal oxidases contribute to the changes in respiratory activity that occur during biofilm growth under light/dark cycling conditions. These findings highlight the strong potential influence of fluctuating environmental and host conditions during infection, and the importance of considering these factors when designing treatment approaches.

## RESULTS

### *Growth with constant light exposure affects TTC reduction activity in* Pseudomonas aeruginosa *PA14 biofilms*

The formation of *P. aeruginosa* biofilms, which is dependent on the extent and timing of biofilm matrix production, is highly sensitive to the availability of respiratory substrates and the capacity for aerobic and anaerobic metabolisms (31, 32). Recently, we showed that *P. aeruginosa* biofilm development is also affected by light exposure and that, interestingly, one of the proteins that regulates this process appears to integrate light and redox information (33). These observations raised the question of whether light exposure affects redox metabolism in *P. aeruginosa*. To test this, we adapted an assay based on reduction of the redox indicator triphenyl tetrazolium chloride (TTC) (34–37). TTC is colorless in its oxidized form; it is reduced by components of the electron transport chain and subsequently forms a red precipitate (**Figure 1A, Figure S1A**). We added TTC to agar-solidified media for the growth of colony biofilms of *P. aeruginosa* PA14 (**Figure 1B**). For these experiments, we used a PA14 Δ*phz* mutant because this strain is unable to produce phenazines, endogenous redox-cycling compounds that have the potential to react with tetrazolium salts and therefore could interfere with the TTC assay (38).

**Figure 1.**
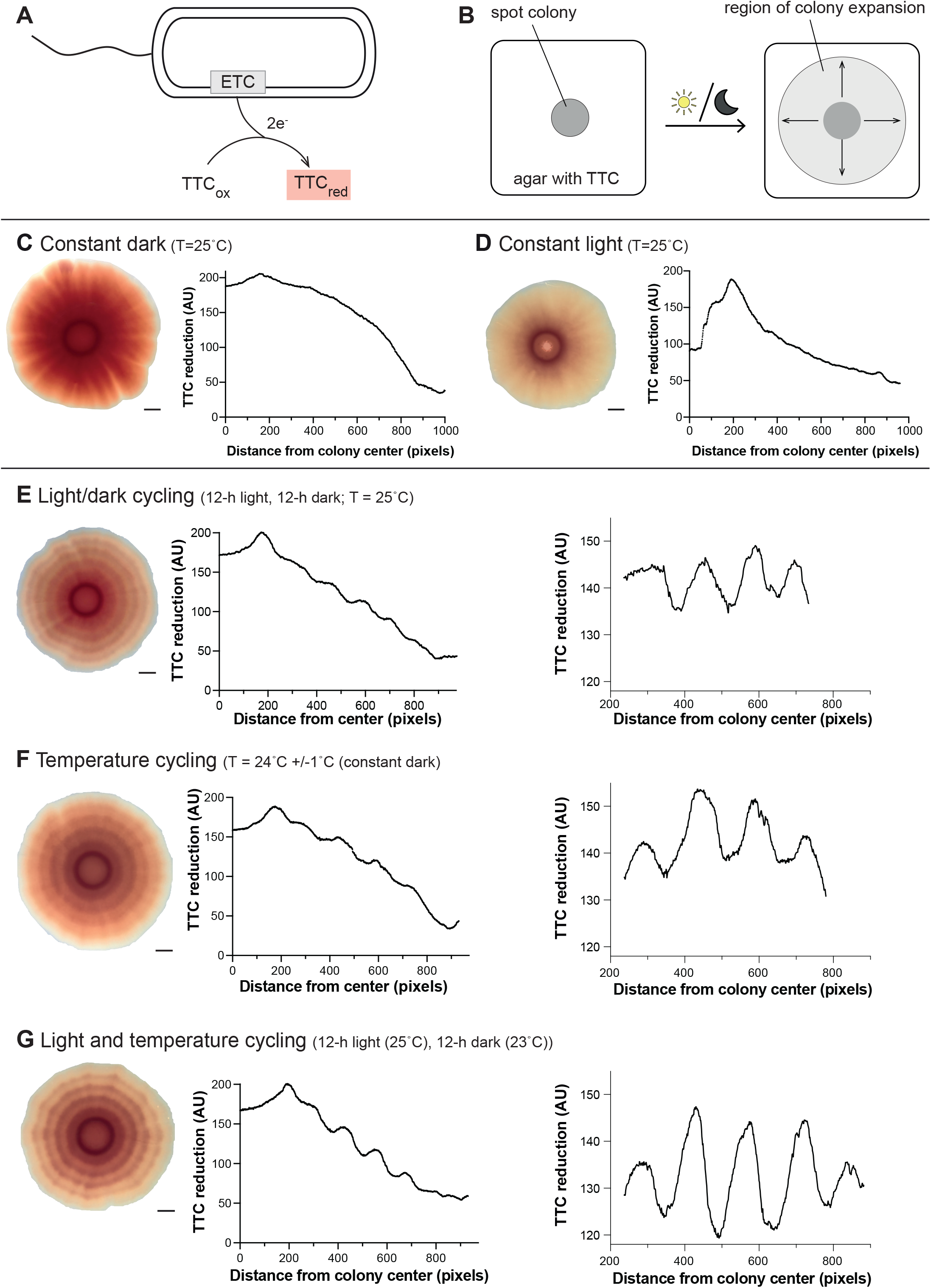
*Pseudomonas aeruginosa* Δ*phz* biofilms grown under light/dark or temperature cycling conditions form patterns of TTC reduction. **(A)** Schematic showing PA14 electron transport chain (ETC)-dependent reduction of TTC, which produces a red precipitate. **(B)** Schematic of experimental setup examining the effect of biofilm growth under constant or cycling conditions (with light or dark conditions indicated by the sun and moon). **(C&D)** Left: Δ*phz* biofilm grown under the indicated condition. Right: Quantification of red color intensity (i.e., TTC reduction) at the indicated distance from the biofilm center for a radius of the biofilm. **(E-G)** Left: Δ*phz* biofilm grown under the indicated conditions. Middle: Quantification of red color intensity (i.e., TTC reduction) at the indicated distance from the biofilm center for a radius of the biofilm. Right: The linear portion of the data was detrended using the linear detrend function of BioDare2 (133). For all experiments, the concentration of TTC in the growth medium was 0.004%. Experiments were performed in biological triplicate and representative data are shown. Scale bars are 2.5 mm.

We grew biofilms on TTC-containing media and compared those grown in darkness to those grown with constant exposure to white light, at an intensity comparable to sunlight on a winter day in the Northern Hemisphere (100 μmol photons m^-2^ s^-1^) (39). We found that growth with constant light led to lower levels of TTC reduction overall than growth in darkness (**Figure 1C, D**). Quantification of colony-forming units for biofilms grown in each of these conditions yielded similar values (**Figure S2E**), indicating that differences in TTC reduction are not due to differences in growth. Biofilms grown in either condition also showed overall decreases in the amount of TTC reduction as they grew, with the biofilm center (older growth) showing more TTC reduction than the biofilm edge (younger growth).

### Light/dark and temperature cycling elicits bands of entrenched TTC reduction activity in PA14 biofilms

On much of Earth’s surface, sunlight exposure occurs for a limited time interval each day. We therefore asked how periodic exposure to light would affect TTC reduction by growing biofilms under conditions of light/dark cycling. After inoculation, biofilms were subjected to 12 hours of white light, followed by 12 hours of darkness, and this 24-hour cycle was repeated 5 times for a total of 6 days of growth. We expected these conditions to yield intermediate levels of TTC reduction throughout the biofilm, comparable to the average of that observed for constant dark and constant light conditions. However, we instead found that biofilms grown with light/dark cycling showed rings of differential TTC reduction that were not present in biofilms grown under constant conditions (**Figure 1E**).

Daily changes in light exposure can be accompanied by changes in temperature, e.g. for bacteria in shallow water or near the surface in soil (26, 29, 30, 40). In addition, bacteria experience recurring temperature fluctuations when colonizing a host that exhibits circadian rhythms in body temperature, which typically varies by 2° to 3°C (9). To test whether small changes in temperature affect TTC reduction, we grew biofilms with temperature cycling (ΔT = 2°C in constant dark or constant light) and found that this condition also yielded characteristic bands of TTC reduction in biofilms (**Figure 1F, Figure S2**). Bands of TTC reduction formed in biofilms grown under temperature cycling set to a range of temperature values: 24 ± 1°C, 24 ± 2°C, 34 ± 1°C, and 35 ± 2°C (**Figure S3A-E**). Combining both light and temperature cycling (i.e., repeating intervals of 12 hours of white light at 25°C followed by 12 hours of darkness at 23°C) gave rise to the most well-defined TTC ring phenotype (**Figure 1G**). Consistent with this, detrended data show that biofilms grown with both light and temperature cycling show a larger amplitude in TTC reduction value than biofilms grown with single cycling conditions (compare **Figure 1G** to **Figures 1E** and **1F**; replicate data are shown in **Figure S3A**). To examine whether light exposure affected growth under cycling conditions, we measured biofilm height across width in thin sections prepared from biofilms grown with light/dark and temperature cycling. We found that biofilm height was not affected by light/dark cycling (**Figure S4A**).

Next, and to better understand the dynamics of TTC reduction, we built a custom-made set up that allowed us to capture images of growing biofilms every 30 minutes (**Figure S1B**). Using these images, we generated kymographs that visualize temporal changes in TTC reduction for pixels appearing at successive time points at the leading edge of the biofilm, along a selected radius (**Figure 2A**). These kymographs show an increase in red color over time for biomass that became visible under either dark or light conditions. However, they also show that biomass that first became visible under dark conditions reduced more TTC overall than prior growth that first became visible under light conditions, thereby yielding the banding pattern. By correlating TTC reduction with the timing of exposure of new biomass to light and dark, we can see that exposure to light leads to an overall decrease in TTC reduction. We then quantified these changes in pixel intensity by averaging the intensity for pixels at a given circumference of the biofilm and tracking how intensity increased over time for this ring of biomass (**Figure 2B**). This analysis revealed discrete metabolic subzones that formed as the biofilm grew outward: Rings that formed in the dark showed, overall, larger increases in TTC reduction (based on final TTC reduction value at the end of 6 days), over the course of the experiment than rings that formed in the light (**Figure 2B, Figure S4B**). Notably, pixel intensities from a specific dark interval approached similar values (“clustered”) (**Figure 2B, top right**), while pixel intensities from a specific light interval showed a broader range of values due to a stronger attenuation of the signal later in the interval (**Figure 2B, bottom right**). Differences in final TTC reduction values were particularly noticeable after the initial phase of rapid reduction (approximately the first 10-15 hours) that occurred in all subzones (**Figure 2B**).

**Figure 2.**
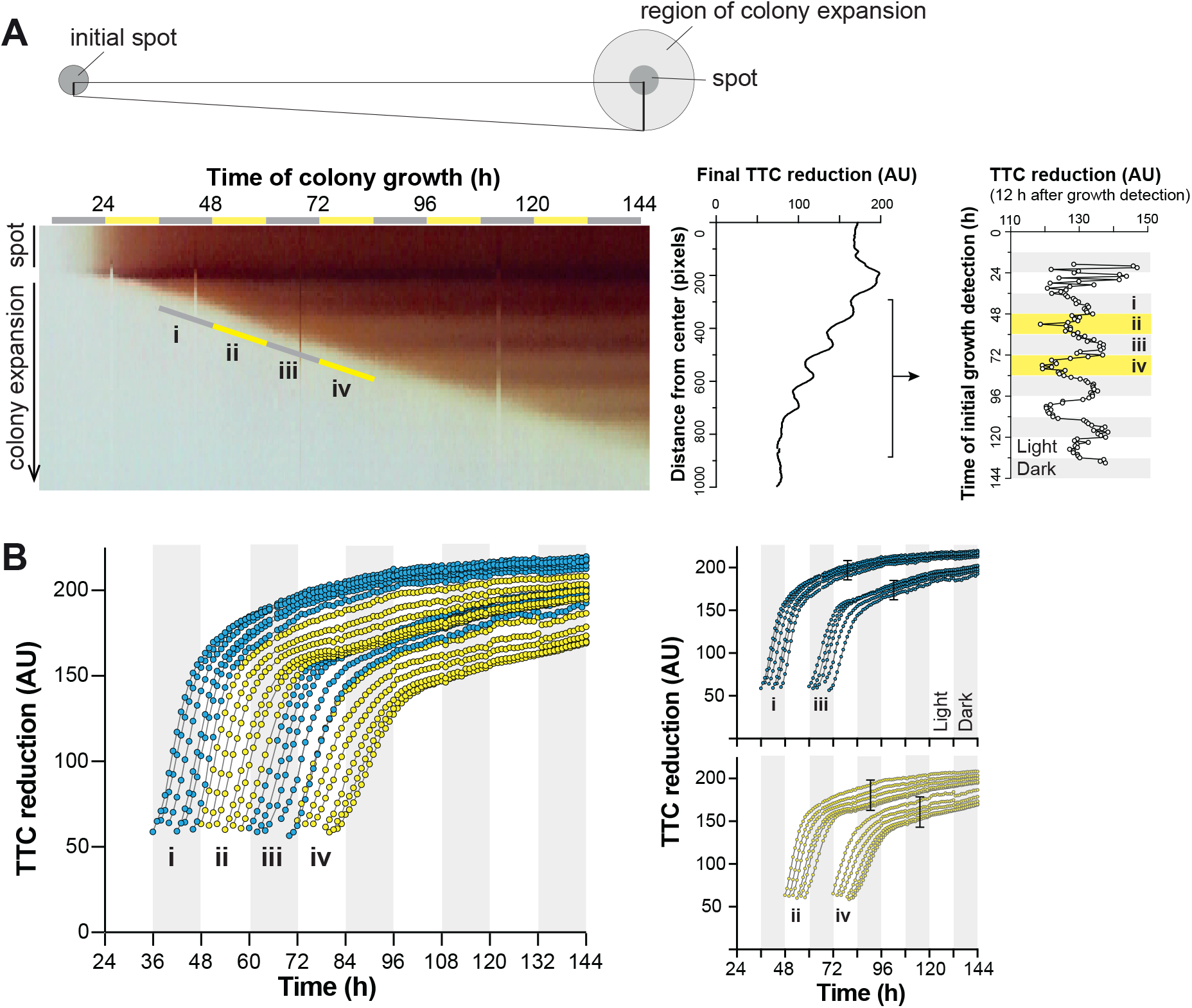
Conditions present during formation of each biofilm growth ring affect the total TTC reduced by the ring. **(A)** Left: Kymograph showing pixel coloration over time across a radius of a biofilm (as depicted in the drawing, top) grown with light/dark and 25°C/23°C cycling conditions on medium containing 0.004% TTC. Bars labeled “i”-”iv” indicate the points at which biomass became detectable via image analysis. Middle: Quantification of red color intensity (i.e., TTC reduction) at the indicated distance from the biofilm center for a radius of the biofilm. Right: Analysis of a time-lapse movie of biofilm growth. The average value of all pixels at the colony edge was tracked over time; the average pixel value, 12 hours after pixels were first detected, is shown. The graph was detrended using the linear detrend function of BioDare2 (133). **(B)** Reduced TTC quantified as average pixel value over time for each ring of growth of the biofilm shown in (A). “i”-”iv” labeling corresponds to pixels of biomass becoming detectable as indicated in (A). Traces corresponding to biomass appearing during dark intervals are shown in shades of blue; those corresponding to biomass appearing during light intervals are shown in shades of yellow. Traces that initiate in dark or light intervals are shown on separate plots, for clarity, on the right. Bars indicate the spread of values for biomass formed in the dark and the light.

Since the combination of light and temperature cycling resulted in the most well-defined TTC reduction phenotype, we used these conditions for additional experiments in biofilms. First, we tested whether the TTC reduction pattern is determined by the duration of intervals during light/dark and temperature cycling, by growing biofilms under shortened (conditions alternating every 6 h) or lengthened (conditions alternating every 24 h) intervals. Time-resolved analysis indicates that interval length dictates changes in TTC reduction; accordingly, relatively low levels of TTC reduction directly correlated with biomass formation in the light (**Figure 3A, B**). Biofilms grown with 6-h intervals showed a lower amplitude of change in TTC reduction value than those grown with 12- or 24-h intervals (**Figure 3B**). Notably, this analysis also revealed that biomass formed in the light shows a “rebound” from decreasing TTC reduction activity at the end of the interval and before the switch to dark conditions, hinting at the presence of a metabolic oscillation. We also tested whether phenazine production affected the TTC ring phenotype by growing PA14 WT biofilms with 12-hour intervals and found that although the rings were indeed less pronounced in this strain (**Figure S2F**), they were nevertheless still detectable and showed similar patterning when compared to Δ*phz*.

**Figure 3.**
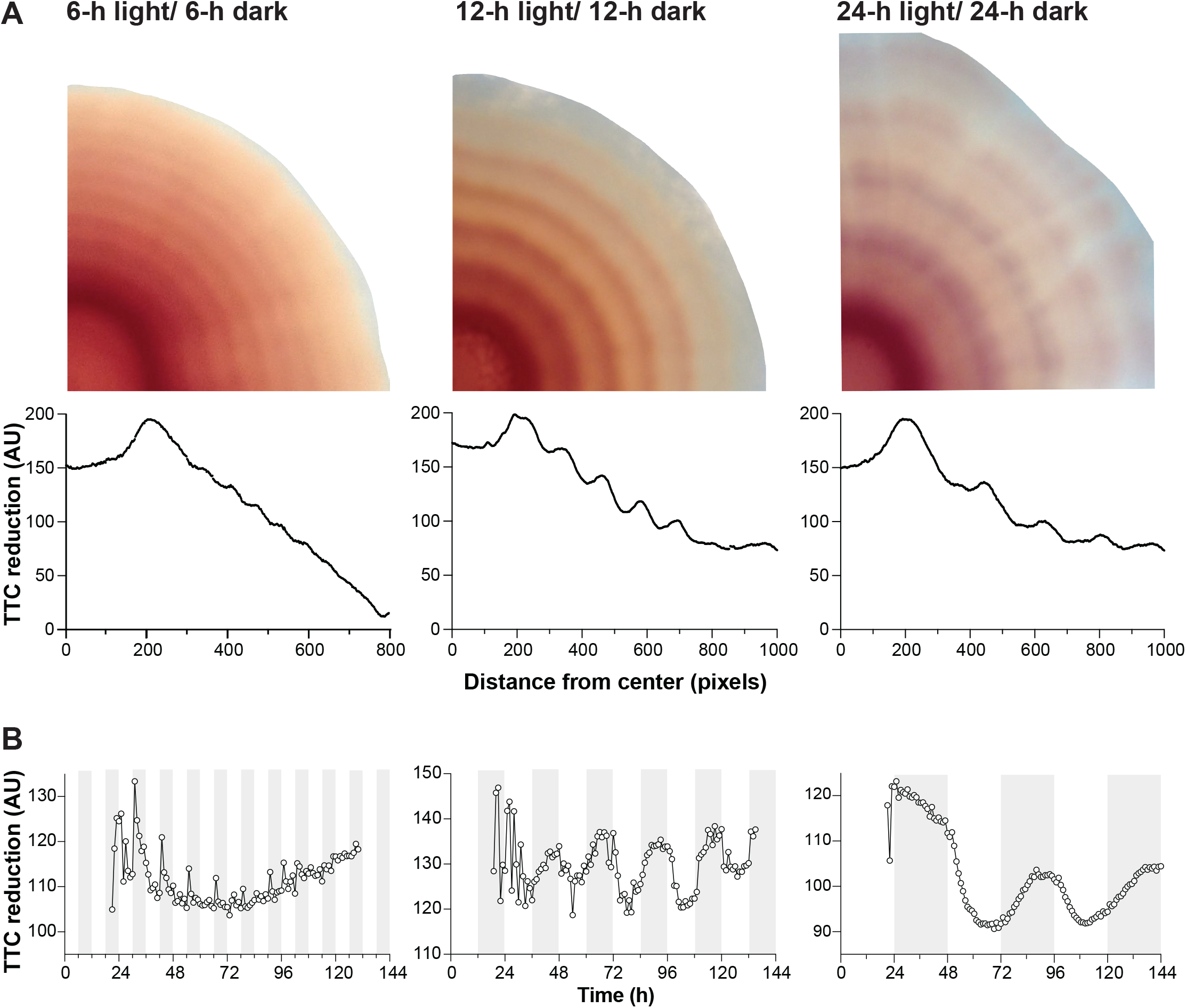
TTC reduction patterns respond to altered periods of light/dark and temperature cycling. **(A)** Top: Biofilms grown under light/dark and 25°C/23°C cycling conditions with 6-hour (left panel), 12-hour (middle panel), or 24-hour (right panel) alternating intervals. Biofilms were grown for a total of 144 hours when using cycling conditions with 6-hour and 12-hour alternating intervals and for a total of 312 hours when using cycling conditions with 24-hour alternating intervals. Bottom: Quantification of red color intensity (i.e., TTC reduction) at the indicated distance from the biofilm center for a radius of the biofilm. The concentration of TTC in the growth medium was 0.004%. Quantification of the final pixel intensity for a radius of each biofilm is shown in the bottom panels. **(B)** Analyses of time-lapse movies of biofilm growth. For each biofilm, the average value of all pixels at the colony edge was tracked over time; the average pixel value, 12 hours after pixels were first detected, is shown. The graph was detrended using the linear detrend function of BioDare2 (133). Experiments were performed with biological triplicates and representative data are shown.

Because an entire biofilm is exposed to cycling conditions, the observed TTC reduction patterns suggest that conditions present during the initial formation of a specific ring of growth serve to “set” the metabolic status of that zone for the remainder of the experiment (**Figure 4A**). To further test whether metabolic states that arise during cycling are maintained in mature biofilms, we repeated the experiment but only exposed the biofilms to TTC after they had experienced the cycling regime. Specifically, we grew colony biofilms under light/dark and temperature cycling conditions for 144 hours on a medium without TTC, then transferred them to a medium containing the redox dye and incubated them for 270 minutes in darkness at 23°C. Indeed, these biofilms formed distinct rings of TTC reduction, particularly visible closer to the biofilm edge (**Figure 4B**), indicating that metabolic status becomes entrenched during initial biofilm zone formation and is then maintained for hours to days.

**Figure 4.**
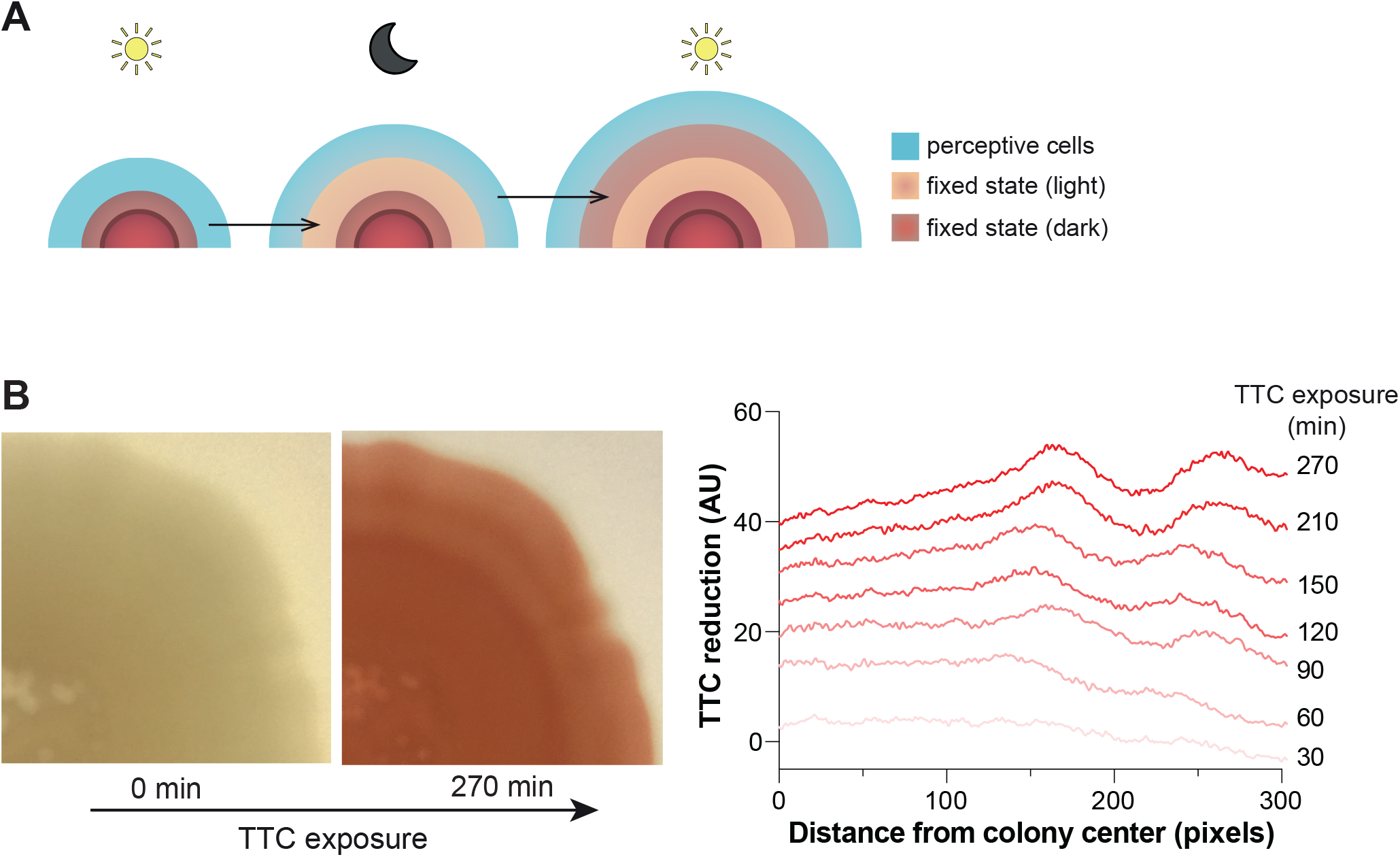
Fixed physiological states are maintained after growth in cycling conditions. **(A)** Model of biofilm development under cycling conditions: The environmental condition (e.g., light or dark) experienced by perceptive cells at the leading edge of the biofilm (blue) “sets” their physiological status, which then remains fixed through subsequent changes in conditions. **(B)** Left panels: Biofilm coloration after 144 hours of growth under light/dark and 25°C/23°C cycling conditions on medium without TTC (“0 min”), and then after transfer to medium containing 0.005% TTC and incubation under constant dark and 23°C conditions for 270 min. Right panel: Quantification of pixel intensity across a selected radius, at the indicated time points after TTC exposure, for the biofilm shown on the left. These TTC reduction values were corrected using the TTC quantification of the pre-transfer biofilm (“0 min”).

### The effect of light/dark cycling on TTC reduction is observed at different wavelengths of light and involves wavelength-specific sensory proteins

While white light comprises a range of wavelengths from ~400-700 nm, physiological responses to light are typically specific to a narrower range of wavelengths due to the unique sensitivities of protein cofactors or small molecules. To test if specific wavelengths are responsible for the effects of light/dark cycling on TTC reduction in biofilms, we grew biofilms under cycling with blue, red, or far-red light while keeping the temperature constant at 25°C. We found that all three conditions yielded rings of TTC reduction like those observed under cycling with white light, indicating that light/dark cycling with different wavelengths of light can affect respiratory electron transport (**Figure 5B**).

**Figure 5.**
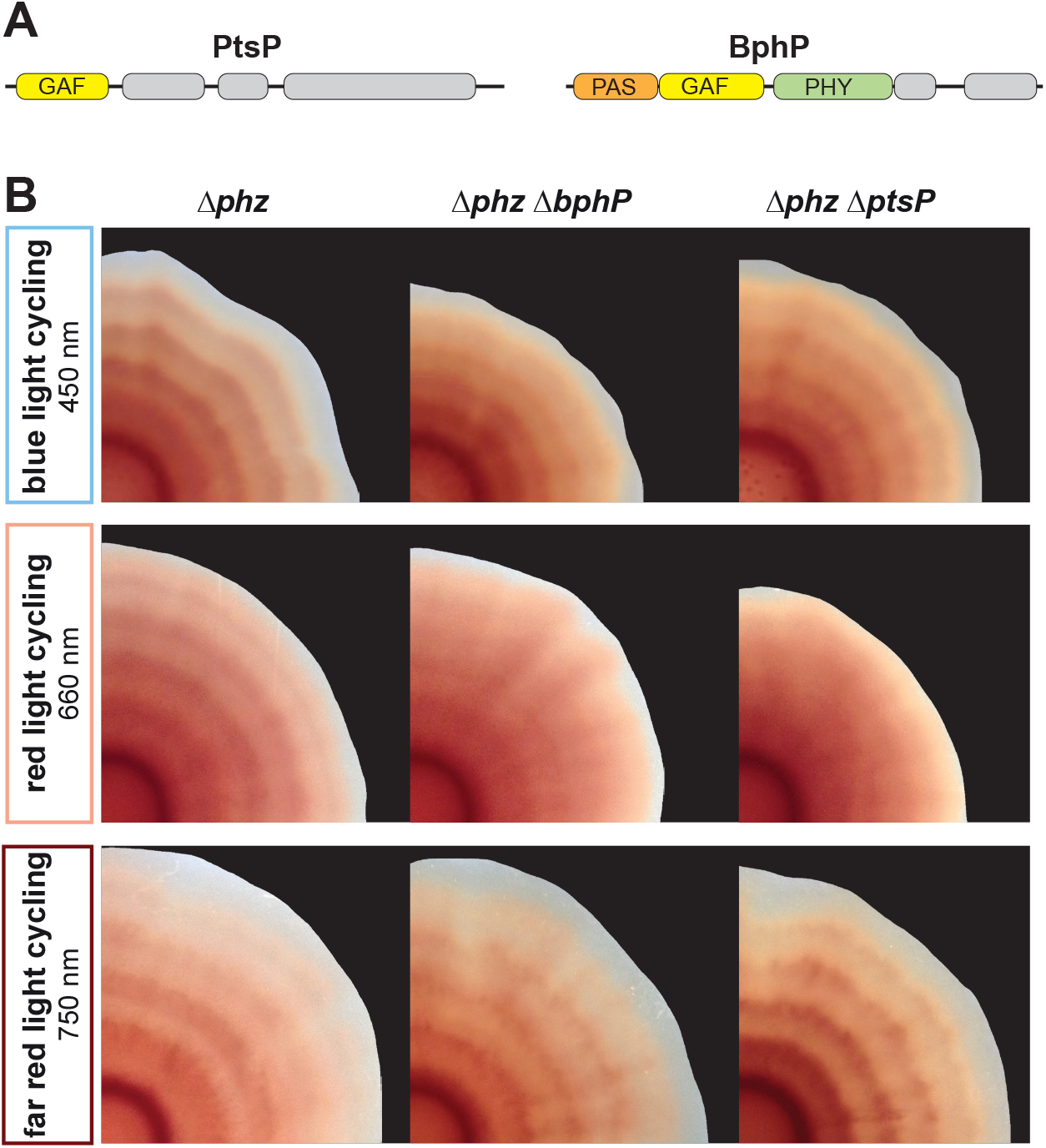
Biofilms grown under light/dark cycling conditions form TTC reduction patterns in blue, red, or far-red light, and Δ*bphP* and Δ*ptsP* mutants show defects in TTC ring pattern formation, specifically under red light. **(A)** Schematics illustrating domain architectures of PtsP and BphP, highlighting putative light-sensing domains. (**B**) TTC reduction patterns in biofilms of the parent strain (Δ*phz*) and indicated mutants grown under light/dark cycling of blue, red, or far-red light with 12-hour intervals. Temperature was kept constant at 25°C and the growth medium contained 0.004% TTC. Experiments were performed with biological triplicates and representative data are shown.

We next asked if PA14 has dedicated sensors that regulate TTC reduction during exposure to each range of wavelengths. Light-sensing proteins have been found in diverse eukaryotes, archaea, and bacteria, including many non-phototrophs (13, 41–44). These proteins contain light-sensitive domains linked to regulatory domains that (i) themselves control intracellular signaling or gene expression, or (ii) interact with other proteins that perform these functions (13, 45, 46). Though the PA14 genome harbors only one gene encoding a canonical light-sensing domain--for the bacteriophytochrome BphP--it also contains genes for dozens of other proteins with PAS or GAF domains, which are broadly implicated in responses to light. These include the protein RmcA, which contains 4 PAS domains, and which mediates an effect of constant light exposure on PA14 biofilm morphogenesis (33). We obtained mutants lacking functional versions of each of the PAS- and/or GAF-domain-containing proteins from the PA14 ordered transposon insertion mutant library (47) and screened for effects on the characteristic TTC reduction pattern that we observe for the parent strain (PA14 WT) under light/dark cycling with blue, red, or far-red light (**Table S1A**). After this initial screen, we generated clean deletion mutants in the *Δphz* background for candidate genes. We found that mutants with disruptions in either *bphP* or *ptsP,* when compared to corresponding parent strains, showed decreased TTC reduction patterning under red light/dark cycling but retained their abilities to produce more pronounced TTC reduction patterns under blue or far-red light/dark cycling (**Figure 5B**). Notably, the mutants lacking functional BphP or PtsP formed biofilms that were similar to the parent strain when grown with temperature cycling under constant dark conditions (**Figure S3F**). Together, these results suggest that BphP and PtsP contribute specifically and in a temperature-independent manner to the effects of red light on TTC reduction in *P. aeruginosa* biofilms. We did not identify candidate blue-light or far-red light sensors in the screen; this indicates that multiple sensors mediate the effects of blue and far-red light on TTC reduction. Blue light and far-red light may also elicit cellular responses because they are directly absorbed by various cellular cofactors and metabolites, including cytochromes, flavins, and by-products of heme degradation (21, 48–51)(52—58).

### Light/dark and temperature cycling yields zones of biofilm growth that are transcriptionally entrenched

As described above, the patterns in biofilms grown under light/dark cycling suggest that the conditions present during the initial formation of a ring of growth serve to determine the metabolic status of that ring for the remainder of the experiment, even though conditions continue to cycle throughout (**Figure 4A**). We hypothesized that these entrenched physiological states would be detectable at the level of gene expression. To test this, we used laser capture microdissection microscopy to sample from four adjacent zones of duplicate biofilms, i.e., two zones formed during light intervals and two zones formed during dark intervals for each of two biofilms and performed RNA sequencing analysis (RNAseq) (**Figure 6A, top**). Principal component analysis confirmed for all but one sample that initial exposure to light or dark resulted in and contributed the most to differential transcriptomic profiles (**Figure S5A**). The outlier sample was excluded from further analysis. We found that although transcriptomes were assessed at a final time point and the entire biofilm had been exposed to light/dark cycling conditions for days, differences in expression were maintained (**Figure 6A bottom, 6B**). This confirms that, as suggested above by our observations of respiratory activity (**Figures 1-3 and 4B**), cells become “physiologically imprinted” by the conditions under which their zone of the biofilm forms. In other words, cells that are situated at the leading edge of the biofilm under dark conditions become strong TTC reducers and express a distinct set of genes from cells that are situated at the leading edge under light conditions, which become weak TTC reducers. We note that principal component PC2 may reflect the difference in age between the two sampled “Dark” zones, i.e., zones formed during the dark intervals, while such a separation between the two sampled “Light” zones was not seen using the first two principal components (**Figure S5A**). Using a 2-fold change cutoff, we found that 341 genes were upregulated in biofilm zones that formed during light exposure and that 170 genes were upregulated in zones formed during dark intervals (**Figure 6B; Supplemental file 1**).

**Figure 6.**
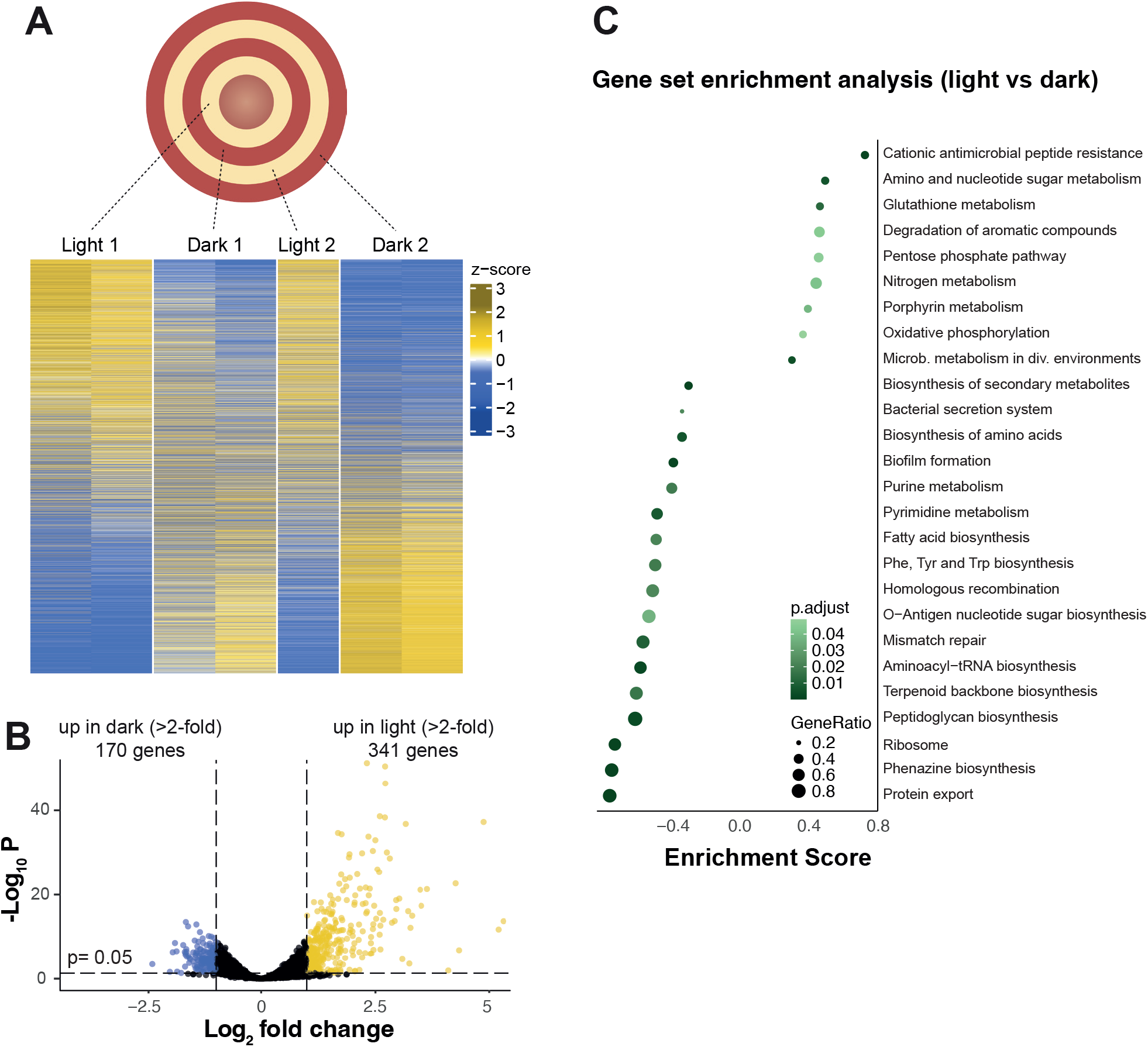
Biofilms grown with light/dark and temperature cycling show fixed physiological responses, evident as differences in transcription in bands of TTC reduction. **(A)** Top: Schematic showing sampling sites in a biofilm grown under cycling conditions on medium containing TTC. The indicated biofilm zones were sampled from duplicate biofilms and subjected to RNAseq. Bottom: Heatmaps representing relative transcript levels (z-scores), shown for each replicate biofilm sample in order of fold change. One sample was omitted because it was determined to be an outlier by principal component analysis. **(B)** Volcano plot showing log_2_-transformed average fold change versus statistical significance for each gene, based on normalized transcript levels detected by RNAseq in biofilm zones formed during light vs. dark intervals. Genes that showed a >2-fold increase in expression under dark/23°C or light/25°C conditions are represented in blue or yellow, respectively. A total of 5945 genes are represented. **(C)** Gene set enrichment analysis using groups of genes based on KEGG-annotated metabolic pathways. Significant enrichment scores above 0 correspond to over-represented groups of genes in biofilm zones from light intervals, while significant enrichment scores below 0 correspond to over-represented groups of genes in biofilm zones from dark intervals. Gene ratio corresponds to the fraction of genes per pathway contributing to the enrichment.

Upon examining the identities of genes affected by light/dark cycling, we noticed a strong representation of those involved in central metabolism, respiration, and redox homeostasis--consistent with the effects of light/dark cycling on TTC reduction. In particular, the KEGG-defined pathways of the pentose phosphate pathway, oxidative phosphorylation, glutathione metabolism, and purine metabolism (59) showed substantial representation of genes that were differentially expressed during light/dark cycling (**Figure 6C**). For each of these pathways, the genes that were differentially expressed were not consistently associated with biofilm zone formation during either light or dark intervals; rather, each pathway contained distinct subsets of genes that were upregulated in zones formed in either the light or the dark. For some pathways, these subsets of genes were linked by common features or functions. In the pentose phosphate pathway, for example, genes involved in the oxidative branch were upregulated in zones formed during light intervals, and genes involved in the non-oxidative branch were upregulated in zones formed during dark intervals (**Figure S5B, C**). For oxidative phosphorylation, we saw two distinct groups of genes for terminal oxidase components that clustered with respect to their expression profiles (**Figure 7A, B**). Genes coding for components of the cyanide-insensitive oxidase (“Cio”), the *bo*_3_ terminal oxidase (“Cyo”), and the *aa*_3_ terminal oxidase (“Cox”) form the first group and were downregulated in rings of growth that formed during dark intervals and upregulated in those that formed during light intervals (**Figure 7A**). In contrast, genes coding for components of *cbb*_3_-type cytochrome *c* oxidase complex CcoN1O1Q1P1 (“Cco1”) and for the orphan terminal oxidase proteins CcoN4Q4, which form the second group, were downregulated in rings of growth that formed during light intervals and upregulated in those that formed during dark intervals (i.e., the opposite expression profile of the first group) (**Figure 7A**). While the RNAseq sampling did not allow us to address terminal oxidase distribution across the depth of the biofilm, for which we have previously demonstrated a differential expression pattern (32, 60), the expression profiles of terminal oxidase genes observed under light/dark and temperature cycling in biofilms nevertheless suggests that the preferential use of terminal oxidases shifts during cycling. We were intrigued to see that the set of terminal oxidase genes showing the largest change in expression during light/dark and temperature cycling was the *cyoABCDE* operon, because this locus is not expressed at high levels in typical aerobic or microaerobic planktonic cultures grown in the laboratory (61). Overall, the effects of light/dark cycling on gene expression in PA14 biofilms are consistent with observations of daily changes in light exposure affecting central metabolism, respiration, and redox homeostasis in other organisms (23, 62–64).

**Figure 7.**
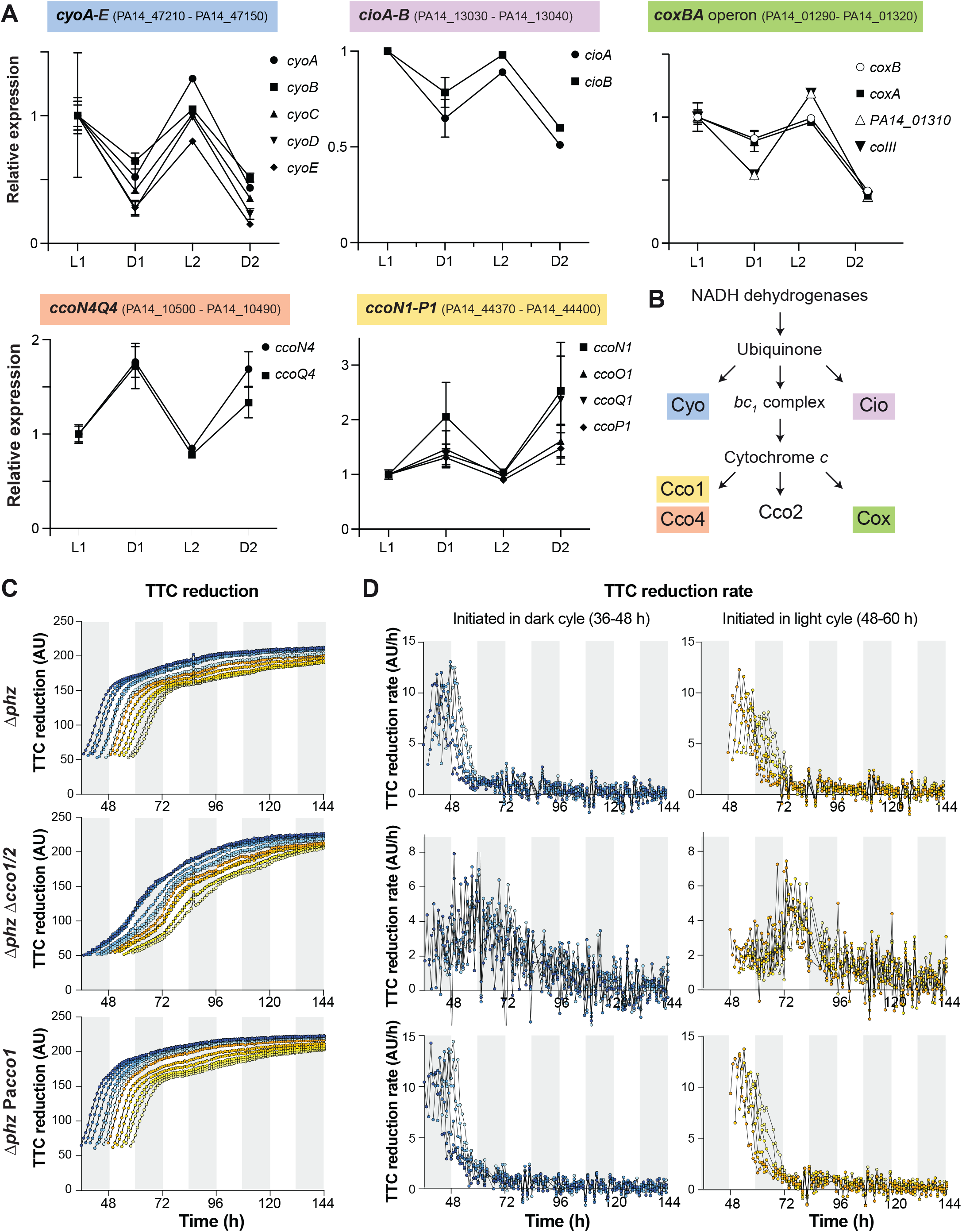
Light/dark and temperature cycling affects the expression of PA14 terminal oxidase genes and *cbb*_3_-type oxidases influence the dynamics of biofilm TTC reduction under this condition. **(A)** Average relative expression of representative genes that code for subunits of terminal oxidase complexes (source: RNAseq analysis in **Figure 6**). “L1” refers to the “Light 1” ring depicted in Figure 6A (Top), “D1” refers to the “Dark 1” ring, and so on. Error bars indicate the standard deviation of duplicate samples. **(B)** Schematic of *P. aeruginosa*’s branched ETC. Cyo, Cio, Cox, Cco2, and Cco1 are terminal oxidase complexes and Cco4 refers to two “orphan” terminal oxidase subunits that can replace the corresponding subunits in Cco2 or Cco1 (32, 65). **(C)** Pixel intensity over time for selected pixels in biofilms of the indicated strains grown under light/dark and temperature cycling on medium containing 0.004% TTC. **(D)** TTC reduction rate over time for data shown in (C), with data for biomass that formed under dark conditions on the left and data for biomass that formed under light conditions on the right. For (C) and (D), data points for biomass formed under dark conditions are blue and those for biomass formed under light conditions are yellow. Darker shades of blue and yellow are used for biomass that formed at earlier time points within an interval and lighter shades are used for biomass that formed at later time points. Results are representative of 3 biological replicates.

### cbb_3_-type terminal oxidases contribute to TTC reduction in rings formed during dark intervals

Components of *cbb_3_*-type terminal oxidases are encoded by 4 different loci in the PA14 genome. Two of these gene clusters, *ccoN1O1Q1P1 (cco1)* and *ccoN2O2Q2P2 (cco2)*, are each sufficient to produce a complete functional oxidase, i.e., they contain all the components of a *cbb_3_*-type complex. Two others, *ccoN3Q3* and *ccoN4Q4,* are “orphan” clusters that each contain two subunits that can substitute for subunits encoded by the 4-gene operons. Genetic and biochemical analyses have shown that *cbb_3_-*type oxidase isomers containing almost all combinations of subunits from each of these loci are functionally similar (65), though the differential expression of *cco* gene loci can affect their relative contributions under specific growth conditions (32, 61). In our RNAseq analysis, we observed that *cco1* and *ccoN4Q4* showed a similar pattern of relatively high expression in biofilm zones that formed during dark intervals and relatively low expression in biofilm zones that formed during light intervals. We therefore hypothesized that *cbb*_3_-type terminal oxidases contribute to TTC reduction in rings formed during dark intervals. Because *P. aeruginosa* possesses multiple terminal oxidases that could function in a compensatory manner (66), we were unsure whether a *cbb_3_*-type terminal oxidase mutant would show a detectable phenotype for TTC reduction. Nevertheless, we grew (i) a Δ*phz*Δ*cco1cco2* mutant, which is unable to produce functional *cbb_3_*-type terminal oxidases and (ii) Δ*phz* PaCco1, which contains Cco1 and the two orphan clusters CcoN3Q3 and CcoN4Q4, but lacks Cio, Cox, Cyo and Cco2, and compared these to the Δ*phz* parent strain on solidified media containing TTC with light/dark and temperature cycling. We then analyzed the increase in pixel intensity over time for each zone of the biofilm as described above for **Figure 3**. TTC reduction curves for biomass that formed in the first 60 hours of growth are shown in **Figure 7C**, and images of corresponding biofilms are shown in **Figure S6**. Similar final TTC reduction values were observed for the biomass formed by Δ*phz*, Δ*phz*Δ*cco1cco2*, and Δ*phz* PaCco1 in the first 60 hours of growth. The Δ*phz* PaCco1 strain and the parent strain also showed similar TTC reduction dynamics, indicating that Cco1 is sufficient to support wild-type patterns of respiration in light/dark cycling conditions. By contrast, the Δ*phz*Δ*cco1cco2* mutant showed altered TTC reduction dynamics, with rates of reduction that were lower than those of the parent strain for most of the incubation (**Figure 7D**). Comparing the Δ*phz*Δ*cco1cco2* mutant to the parent strain, we observed that the increase in TTC reduction rate was more gradual and that maximum rates of reduction tended to occur ~10-20 hours later, i.e., roughly one interval later, for the mutant lacking *cbb*_3_-type oxidases.

## DISCUSSION

Though chemotrophic bacteria do not depend on light as an energy source, they are impacted by light exposure, or fluctuations in light exposure, in several ways. Light affects environmental conditions by driving the metabolisms of phototrophs, which produce oxidants and other resources that can be used by chemotrophs (67–69). Light can also directly damage cofactors required for chemotrophic metabolisms (21, 48–51) and increase the temperature of exposed soil, sediment, or water (26, 30, 40, 70). Finally, light acts as a cue to entrain endogenous oscillations in the physiologies of circadian organisms, such as fluctuations in body temperature and elements of the immune response, e.g. phagocytic activity (71, 72). Bacteria that colonize circadian organisms are therefore indirectly affected by the entraining effects of light on their hosts and the host environment (9).

We investigated whether growth under light/dark and temperature cycling affected metabolism and gene expression in colony biofilms of *P. aeruginosa* PA14, a chemotrophic bacterium and opportunistic pathogen. We found that growth of PA14 biofilms under either light/dark or temperature cycling promoted the formation of rings of alternating levels of metabolic activity, as indicated by reduction of the dye TTC. When biofilms were grown under conditions of alternating intervals of dark at 23 °C and light at 25 °C, the formation of rings with increased activity correlated with growth under dark and low-temperature conditions and adjusted to the period length. We also found that cycling of blue, red, and far-red light were each sufficient to elicit ring formation. RNAseq analysis of samples taken from rings of high or low TTC reduction activity revealed differences in the expression of genes involved in primary metabolism and redox homeostasis. Our results suggest that the conditions that prevail during growth of a specific zone of the biofilm serve to “set” the physiological status of that zone, because although entire biofilms were exposed to cycling conditions, individual zones of these biofilms showed sustained differences in TTC reduction activity and gene expression. Consistent with this, we also observed the formation of bands of TTC reduction in biofilms that were grown under cycling conditions on a medium without the dye and subsequently transferred to a medium containing the dye.

Ring or band patterning has been described for expanding multicellular structures formed by various chemotrophic bacteria and fungi. Perhaps the best-known of these is the model microbial fungus *Neurospora crassa.* Its spatial conidiation pattern is used as a readout for circadian rhythms because conidiation reoccurs daily under specific redox conditions during fungal outgrowth (73). *N. crassa* rhythmic biology has been shown to conform to the definition of a circadian phenomenon and the biochemical basis of circadian regulation in this organism is well understood. Recently, colony biofilms of the soil and human gut commensal bacterium *B. subtilis* were shown to exhibit rings of gene expression that form autonomously (74). Circadian changes in gene expression have also been associated with biofilm formation in this organism (75). Other microbes that have been shown to form ring or band patterns that exhibit rhythmicity include *P. putida*, *L. monocytogenes*, *K. aerogenes*, and dimorphic and cold-adapted yeast (76–80). While some of these phenotypes require growth under light/dark or temperature cycling conditions, others occur autonomously.

We observed that light/dark cycling during growth of *P. aeruginosa* biofilms affected respiratory activity and redox metabolism. Control of primary metabolism and redox homeostasis in response to light/dark cycling or as part of a biological oscillation has been reported for diverse phototrophs (as would be expected for organisms that use light as an energy source; (81–83)) and non-phototrophs (62, 84, 85). A comprehensive study of organisms representing all domains of life showed circadian oscillations between the oxidized and superoxidized states of peroxiredoxins, broadly conserved enzymes that control intracellular levels of reactive peroxides and that can serve as indicators of redox homeostasis. These previous findings suggest that circadian rhythms evolved in association with daily recurring changes in cellular redox metabolism (23). In the budding yeast *Saccharomyces cerevisiae*, two types of respiratory oscillations have been observed in continuous liquid cultures: a circadian oscillation that is dependent on temperature cycling over a 24-hour period (86), and an autonomous, ultradian phenomenon referred to as the yeast respiratory oscillation or the yeast metabolic cycle (YMC). The YMC has been extensively characterized, with multiple studies describing the changes in central metabolism and respiration that occur over the course of the cycle (24, 84, 87, 88), and recent evidence suggests that this temporal organization of metabolism optimizes cellular processes in a manner that confers stress resistance (89). Among genes that show the highest periodicity in the YMC are COX10, involved in heme *a* biosynthesis (a pathway that converts heme *b* to heme *a* via a stable heme *o* intermediate) (84, 90), and PET117, involved in assembly of the mitochondrial (*aa*_3_-type) terminal oxidase (91). Interestingly, the *cyo* operon, which includes the heme *o/a* biosynthetic gene *cyoE,* was upregulated in zones that formed at the colony edge during light intervals. The *P. aeruginosa aa*_3_-type oxidase also showed periodicity and upregulation in biomass that forms in the light under cycling conditions. Thus, in both yeast and *P. aeruginosa*, periodicity of these genes during respiratory oscillations and growth in cycling conditions likely reflect increased demand for the terminal oxidases that use *a*- and *o-*type hemes, i.e., Cox/mitochondrial terminal oxidase and the Cyo terminal oxidase.

Microbial physiology can be sensitized to light exposure when (i) light damages cofactors that are required for protein function (20, 48, 49, 51, 92, 93), or (ii) when light stimulates a conformational change in a protein that participates in regulatory interactions (15, 82, 94, 95). YMCs occur in the absence of light/dark cycling but show altered period and amplitude during constant exposure to blue light. These effects appear to arise from light-dependent inhibition of respiratory cytochromes, which show absorption peaks that correlate with wavelengths that have the greatest impacts on the oscillations (21). While a similar inhibition could contribute to the fluctuation in respiratory activity reported here, which occurs in *P. aeruginosa* biofilms grown under light/dark cycling, our observations indicate that it is not the sole elicitor of this pattern. First, the formation of rings of reduced TTC occurs in *P. aeruginosa* biofilms grown in cycling conditions with blue, red, or far-red light and is not specific to wavelengths that are maximally absorbed by cytochromes. Second, we identified proteins with potential regulatory functions that contribute to ring formation. The first hit, BphP, was a light-sensing histidine kinase that contains the N-terminal phytochrome domain architecture characteristic of these proteins (**Figure 5A**). *P. aeruginosa* BphP has previously been shown to affect expression of a regulon that includes genes for respiratory complexes and the synthesis of carbon storage compounds (96); in the YMC, these carbon storage compounds are hypothesized to act as electron donors during the high-respiration phase of the YMC. BphP has also been shown to act on the response regulator AlgB to affect regulatory networks and biofilm development in response to light (13, 33). The second screen hit, PtsP, was a phosphotransferase that contains an N-terminal GAF domain (**Figure 5A**). GAF domains are common sensory domains that can detect a broad range of stimuli and that are associated with light sensing in proteins from diverse organisms. PtsP is the first component of the phosphoryl transfer system (“Pts”), a signaling cascade that controls diverse processes, including aspects of central metabolism, in pseudomonads and other bacteria (97). Interestingly, it has been suggested for *Acinetobacter baumannii* that the target of PtsN, PtsO, acts as an anti-sigma factor for RpoE (also known as AlgU or AlgT), which is part of the AlgB-dependent regulatory network (98, 99)--hinting that BphP and PtsP may constitute independent sensors with converging downstream effects. Notably, although the protein RmcA makes a strong contribution to morphological changes in light-exposed biofilms (14), we found that the *ΔphzΔrmcA* mutant showed a TTC reduction phenotype similar to that of the parent strain when grown under light/dark cycling conditions. Overall, the results of our study are consistent with a model in which light affects respiratory activity both through effects of specific wavelengths on cytochromes in the electron transport chain and effects on the conformation of light-sensing regulatory proteins.

Temperature is a key modulator of physiology because it affects membrane fluidity, enzyme activity, and the stabilities of proteins and nucleic acids (100–105). The physiological responses to temperature that have been characterized in bacteria range from (i) transcriptional responses to sudden up- or downshifts in temperature, to (ii) adaptation to a new temperature optimum via prolonged growth in the nonoptimal condition (106–108), to (iii) control of biological thermosensors that affect regulatory pathways and behavior. The first of these constitute the heat and cold shock responses, which serve to counteract harmful effects arising from sudden increases or decreases in temperature (109, 110)(111, 112)(106–108)(100). Examples of adaptation to thermal niche include those found in studies of commensal or pathogenic bacteria, where strains of the same bacterial species have been shown to exhibit differing thermal tolerances depending on the host mammal (113). Bacteria can also become highly specialized to certain temperature regimes, for instance as insect mutualists (114). A diverse array of mechanisms allow bacteria to sense temperature and transduce this information to affect regulation of gene and protein expression and behavior (103). A recent example of this arose in the identification of a thermosensory PAS domain in a protein that modulates *P. aeruginosa*’s cellular levels of c-di-GMP, a messenger molecule with global effects on development and motility (115). Finally, temperature changes can also serve as cues to entrain circadian clock machineries and phenotypic outputs (116–118).

*P. aeruginosa* is typically found associated with eukaryotic organisms or in soil and aquatic environments that have been contaminated by human activity (119). It can colonize a broad range of hosts (120–123). In humans, *P. aeruginosa* can cause a variety of infections, including pneumonia, sepsis, and skin and soft tissue infections (124–126). Depending on the specific body site it is colonizing, *P. aeruginosa* can be directly exposed to light or it can experience the effect of light on host physiology. Light entrains host circadian rhythms, which include fluctuations in body temperature and variations in immune system function (127, 128). By responding to light/dark and temperature cycling, *P. aeruginosa* might have the ability to tune its physiology for optimal survival during association with hosts. We observed that *P. aeruginosa* biofilms exhibit metabolic switching at temperatures comparable to those encountered on human skin (33°C - 37°C) (**Figure S3D, E**). A similar capability has been described for the commensal enteric bacterium *Klebsiella aerogenes*, which shows rhythmic gene expression that is entrained by temperature changes in this range (78, 116, 129). Host settings are unique nutritional environments, and it is now recognized that metabolic adjustments appropriate to these settings are key to pathogen success (130). A better understanding of the effects of light/dark and temperature cycling on pathogen physiology could aid in the development of treatment approaches, which could, for example, be timed to maximize their effects or augmented by modification of light and temperature parameters in situ.

## MATERIALS AND METHODS

### Bacterial culture and light/dark cycling assay

All species and strains used in this study are listed in **Table S1A**. For light/dark (L/D) cycling assays, 1% tryptone (Teknova) plus 1.25% agar (Teknova) was autoclaved and allowed to cool to 60°C and optionally amended with 0.001% to 0.005% 2,3,5-triphenyltetrazolium chloride (TTC) (Sigma Aldrich, Cat # T8877), depending on experimental procedure. Portions (90 ml) of the agar mixture were poured into UV-sterilized, square, gridless plates (Greiner Bio-One, Cat # 688102) and after agar plates solidified they were stored in a dark chamber at 25°C overnight (14 h to 24 h). Bacterial liquid cultures (3 ml of lysogeny broth (LB; (131))) were inoculated using single colonies and shaken at 250 rpm at 37°C overnight (12 h to 16 h). Subcultures were prepared by diluting overnight cultures 1:100 into 3 ml of LB (Δ*phz*Δ*cco12* cultures were diluted 1:50) and grown at 37°C with shaking (250 rpm) until midexponential phase was reached (0.4 to 0.6 AU absorbance measurements at 500 nm, OD_500_). Agar plates were dried in a ventilated, sterile biosafety cabinet for 60 min. One or 5 μl of subculture were spotted onto agar plates. Colonies on agar plates were placed in a temperature-controlled (0.2°C error) incubator with a fitted lighting system (Percival Scientific CU-22LC9). Colony biofilms were grown at 25°C (unless otherwise indicated) at 90-100% humidity for 6 days (or specified if longer) under various lighting conditions (specified below). Images were taken with a Zeiss AxioZoom.V16 Fluorescence zoom Stereomicroscope fitted with an iPod camera or as time-lapse movies using a custom-built movie chamber as described in (33). Time-lapse images were acquired every 30 min. All images were taken in the dark to ensure that the pixel values in the images were consistent.

### Light conditions for light/dark cycling experiments

For most light/dark cycling experiments white light exposure (bulbs: F17T8/TL841/ALTO, Philips) was calibrated to 68.64 μmol photons m^-2^ s^-1^. Light intensities were measured as photon flux density between 400 nm and 700 nm using an AMOUR-SL-125-PAR meter (Biospherical Instruments) for all light bulbs except the far-red light bulbs. For the wavelength-specific light exposure experiments (**Figure 5**), blue light photon flux density (bulbs: ELA-085, F17T8/BL-450, Percival Scientific) was set to 22.1 μmol m^-2^ s^-1^, and red light photon flux density (bulbs: ELA-086, F17T8/IR-660, Percival Scientific) was set to 40 μmol m^-2^ s^-1^. Light intensities for peak wavelengths of all light bulbs were measured using an S130C photodiode power sensor (Thorlabs) with a PM100A power meter console (Thorlabs). For comparison between light bulbs, white light flux density (analyzed at 436, 545, and 612 nm) was measured as 2.66, 1.31, and 1 W/m^2^, respectively. Blue light flux density (analyzed at 422 nm) was measured as 1.11 W/m^2^, red light flux density (analyzed at 660 nm) was measured as 0.87 W/m^2^, and far-red light flux density (analyzed at 750 nm; bulbs: ELA-087, F17T8/IR-750, Percival Scientific) was measured as 0.87 W/m^2^. For the majority of experiments light/dark cycles were set to a 24-hour period (12 hours of light and 12 hours of dark). For experiments with shortened and prolonged light/dark cycles (**Figure 2**), periods were set to 12 hours (6 hours of light at 25°C and 6 hours of dark at 23°C) or 48 hours (24 hours of light at 25°C and 24 hours of dark at 23°C).

### Exposure to TTC post-completion of light/dark cycling assay

For exposure of mature biofilms to TTC (**Figure 4B**), agar plates were prepared as for other light/dark cycling experiments except that the growth medium was not amended with TTC. Bacterial liquid cultures of Δ*phz* Δ*pel* mutants were prepared as described above and spotted on Whatman filter disks. Bacterial colonies were grown under the following cycling conditions: 12-hour white light exposure at 25°C, followed by 12-hour dark at 23°C. Five days after the colonies were spotted, tryptone agar plates were prepared and amended with 0.005% TTC. After 6 days of growth, biofilms were transferred to TTC-amended tryptone agar by carefully lifting each filter disk and placing it on the agar. Images of TTC reduction were taken every 30 or 60 min (final image after 270 min) using the AxioZoom setup as described above.

### Deletion mutants

Markerless gene deletions were created as described in (32). Combinatorial deletion mutants were constructed by successive single deletions of respective genes. Deletion mutants in *P. aeruginosa* PA14 are listed in **Table S1A**. Primers used to amplify 1 kb of flanking sequence from each side of the target gene are listed in **Table S1B**. Plasmids harboring the gene deletion constructs are listed in **Table S1C**.

### RNA sequencing (RNAseq) and transcriptomic analysis

Biofilms were grown in light/dark cycling with 2°C temperature changes (light/25°C vs. dark/23°C in 12-hour intervals) on 0.22 μm polypropylene filters atop agar-solidified medium for 76 hours. The growth medium was 1% tryptone plus 1% agar amended with 0.0025% TTC. Prior to sample collection, each biofilm was placed on a slide with 100 μl RNAlater (Thermo Fisher). Samples were collected by laser capture microdissection microscopy using a Zeiss PALM MicroBeam IV from the outermost four rings (two red rings, i.e., Dark 1 and Dark 2, and the two white rings, i.e., Light 1 and Light 2) for two separate colony biofilms. Areas of the biofilms ranging from 1 to 10 mm^2^ were dissected and transferred to tubes containing 200 μl RNAlater (Thermofisher). Samples were homogenized using a motorized pestle and total RNA was extracted and purified using a RNeasy Plant Mini Kit and RNeasy MinElute Cleanup Kit (Qiagen). RNA was sent to GENEWIZ, Inc. for sequencing with Illumina HiSeq. Reads were mapped using Geneious Prime 2022.0 (https://www.geneious.com) and differential gene expression was determined (method: Geneious; transcripts were normalized by median of gene expression ratios). The heatmap was generated using the R packages ComplexHeatmap, RColorBrewer, and circlize by scaling log-transformed normalized counts by row (gene). Dimension reduction was performed by principal component analysis (depicted in **Figure S5A**) using the R package DESeq2. Significant (adjusted *p*-value < 0.05) and highly differentially expressed (|log_2_FC| > 1) genes were visualized in the volcano plot (**Figure 6B**). Gene expression data were arranged into metabolic pathway groupings using the *P. aeruginosa* UCBPP-PA14 specific entry in the KEGG database (www.kegg.jp) (59) and the R package KEGGREST, and enrichment analysis was performed using the R package clusterProfiler.

### Colony biofilm growth and TTC reduction: movie analysis

All RGB movie images were converted to grayscale. The 2D agar/colony boundary was determined for each image using a thresholding approach: Initially, the final movie image was used to approximate the agar/colony boundary and determine the average pixel intensity value for the entire agar area. Then, a Python script (**Supplemental file 2**), was used to scan from the agar area toward the edge of the colony. The agar/colony boundary was defined by a change in pixel intensity that exceeded three standard deviations from the average agar intensity. The intensity value at the agar/colony boundary in the final image was then used as the threshold to analyze all images and the time at which each pixel gained the threshold intensity was determined. Subsequent intensities were recorded and yielded data that allowed for the generation of TTC reduction graphs (**Figures 3B and 7C**).

### Graphing

TTC reduction values were determined in Fiji image processing software (132). The first and last peaks of TTC reduction were removed before detrending to ensure only the linear section of the graph was further analyzed. Curves were then linearly detrended using the open-access web tool BioDare2 (https://biodare2.ed.ac.uk) (133).

### Thin sectioning analysis

Two layers of 1%tryptone with 1.25% agar were poured to depths of 4.5 mm (bottom) and 1.5 mm (top). Overnight precultures were diluted 1:100 in LB and grown for 2.5 hr, until early-mid exponential phase. 2 μl of subculture were then spotted onto the top agar layer and colonies were grown in 12-h light (25°C), 12-h dark (23°C), and temperature cycling (Percival CU-22L) for 3 days. Bioflms were covered by a 1.5-mm-thick 1% agar layer to prepare for thin sectioning. Colonies sandwiched between two 1.5-mm agar layers were lifted from the bottom layer and fixed in 4% paraformaldehyde and PBS (pH 7.4) overnight at 25°C. Fixed colonies were washed twice in PBS and dehydrated through a series of ethanol washes (25%, 50%, 70%, 95%, 3 × 100% ethanol) for 60 min each. Colonies were cleared via three 60-min incubations in Histoclear-II (National Diagnostics [Atlanta, GA] HS-202) and infiltrated with wax via two separate washes of 100% Paraplast Xtra paraffin wax (Thermo Fisher Scientific 50-276-89) for 2 hr each at 55°C, then colonies were allowed to polymerize overnight at 4°C. Tissue processing was performed using an STP120 Tissue Processor (Thermo Fisher Scientific 813150). Trimmed blocks were sectioned in 10-μm-thick sections perpendicular to the plane of the colony using an automatic microtome (Thermo Fisher Scientific 905200ER), floated onto water at 42°C, and collected onto slides. Slides were air-dried overnight, heat-fixed on a hotplate for 1 hr at 45°C, and rehydrated in the reverse order of processing. Rehydrated colonies were immediately mounted in Prolong Diamond Antifade Mountant (Thermo Fisher Scientific P36961) and overlaid with a coverslip. Differential interference contrast (DIC) images were captured using an LSM800 confocal microscope (Zeiss, Germany). Each strain was prepared in this manner in at least biological triplicates.

## Supporting information

Supplemental Table 1A-C and references

## ACKNOWLEDGEMENTS

This work was supported by NIH/NIAID grant R01AI103369 and NSF award 1553023 to L.D. Colony biofilm samples were collected by laser capture microdissection at the Confocal and Specialized Microscopy Shared Resource of the Herbert Irving Comprehensive Cancer Center at Columbia University, supported by NIH/NCI grant P30CA013696.

## SUPPLEMENTAL MATERIAL

**Figure S1. (A)** Schematic showing the structure and reduction of TTC. **(B)** Schematic of incubation setup used to generate time-lapse movies of colony biofilm growth and TTC reduction.

**Figure S2. (A-C)** Left: Replicates of biofilms grown in constant dark at 25 °C (A), in constant light at 25°C (B), and light and temperature cycling, 12-h light 25 °C, 12-h dark 23 °C (C). Right: Quantification of TTC reduction. **(D)** Comparison of representative TTC reduction quantification from all three conditions in panels (A-C). **(E)** Colony forming units for biofilms grown in constant light or constant dark at 25°C for 4 days. **(E)** Colony forming units for biofilms grown in constant light or constant dark at 25°C for 4 days. **(F)** Top: WT biofilm grown in light and temperature cycling, 12-h light 25 °C, 12-h dark 23°C. Bottom: Quantification of TTC reduction.

**Figure S3. (A)** Left: Representative replicate for Figure 1F. Right: Quantification of red color intensity (i.e., TTC reduction) at the indicated distance from the biofilm center for a radius of the biofilm for three replicates. **(B)** Left: Representative replicate for Δ*phz* biofilm grown in constant light with 24 °C +/- 1°C temperature cycling. Right: Quantification of red color intensity (i.e., TTC reduction) at the indicated distance from the biofilm center for a radius of the biofilm for three replicates. **(C)** Left: Representative WT or Δ*phz* biofilm grown in constant dark with 24°C +/- 2°C temperature cycling. Right: Quantification of TTC reduction for three replicates per condition. **(D&E)** Representative replicates for Δ*phz* biofilms grown in constant dark with 34°C +/- 1°C and 35°C +/- 2°C temperature cycling. **(F)** Light sensor mutants grown in constant dark with 24°C +/- 1°C temperature cycling, on medium containing 0.004% TTC.

**Figure S4. (A)** Representative thin sections of Δ*phz* biofilms grown on medium containing 0.004% TTC for 3 days. Top: Section of biofilm grown with L/D and 24 ± 1 °C cycling. Bottom: Section of biofilm grown in the dark and at 25°C. Scale bars are 200 μm. For each condition, three replicates were quantified and graphed. Biological triplicates are shown.

**(B)** Replicates for figure 2B. Traces corresponding to biomass appearing during dark intervals are shown in shades of blue; those corresponding to biomass appearing during light intervals are shown in shades of yellow.

**Figure S5. (A)** Dimensionality reduction by principal component analysis was applied to RNAseq data for the full set of 8 samples obtained as shown in **Figure 6A**. Ninety percent of maximum possible variance was accounted for by the first two principle components PC1 and PC2. Each sample is plotted according to its values for PC1 and PC2. **(B)** Sixty-seven percent of genes involved in the pentose phosphate pathway were identified in the RNAseq as showing differential expression in biomass formed under light/dark and temperature cycling. Genes coding for enzymes involved in the oxidative branch were upregulated in biomass that formed under light/25°C conditions, and genes coding for enzymes involved in the non-oxidative branch were upregulated in biomass that formed under dark/23°C conditions. The heatmap shows genes involved in the pentose phosphate pathway, arranged according to their relative periodicity in samples from biomass formed under light/dark and temperature cycling. **(C)** Schematic of the pentose phosphate pathway highlighting selected genes whose expression was affected by growth under light/dark and temperature cycling conditions.

**Figure S6.** Images of Δ*phz*, Δ*phz*Δ*cco1cco2*, and Δ*phz* PaCco1 colony biofilms, corresponding to the analyses shown in Figure 7C and D.

**Table S1A. Strains used in this study**

**Table S1B. Primers used in this study**

**Table S1C. Plasmids used in this study**

**Supplemental file 1. List of RNAseq genes**

**Supplemental file 2. Python code for colony time-lapse imaging**

