## Supplemental Figure 1 for "Light/dark and temperature cycling modulate metabolic electron flow in *Pseudomonas aeruginosa* biofilms"

**A**

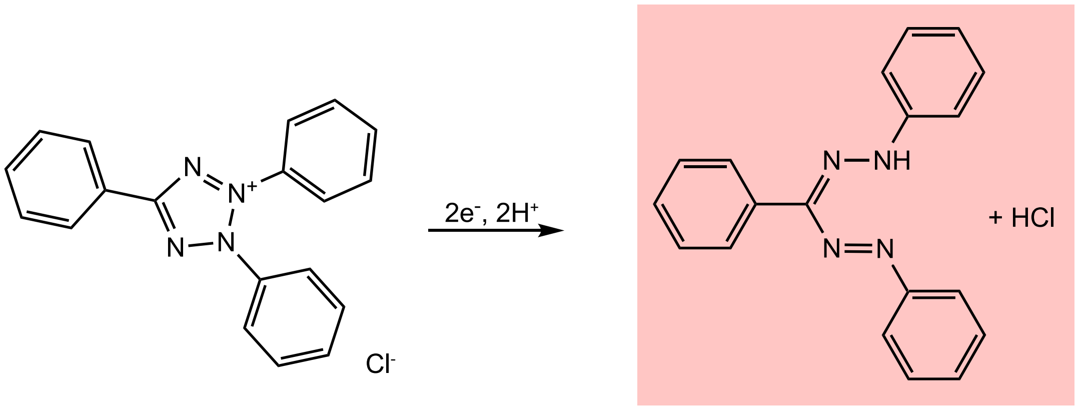


2,3,5-triphenyltetrazolium chloride

1,3,5-triphenylformazan

**B**

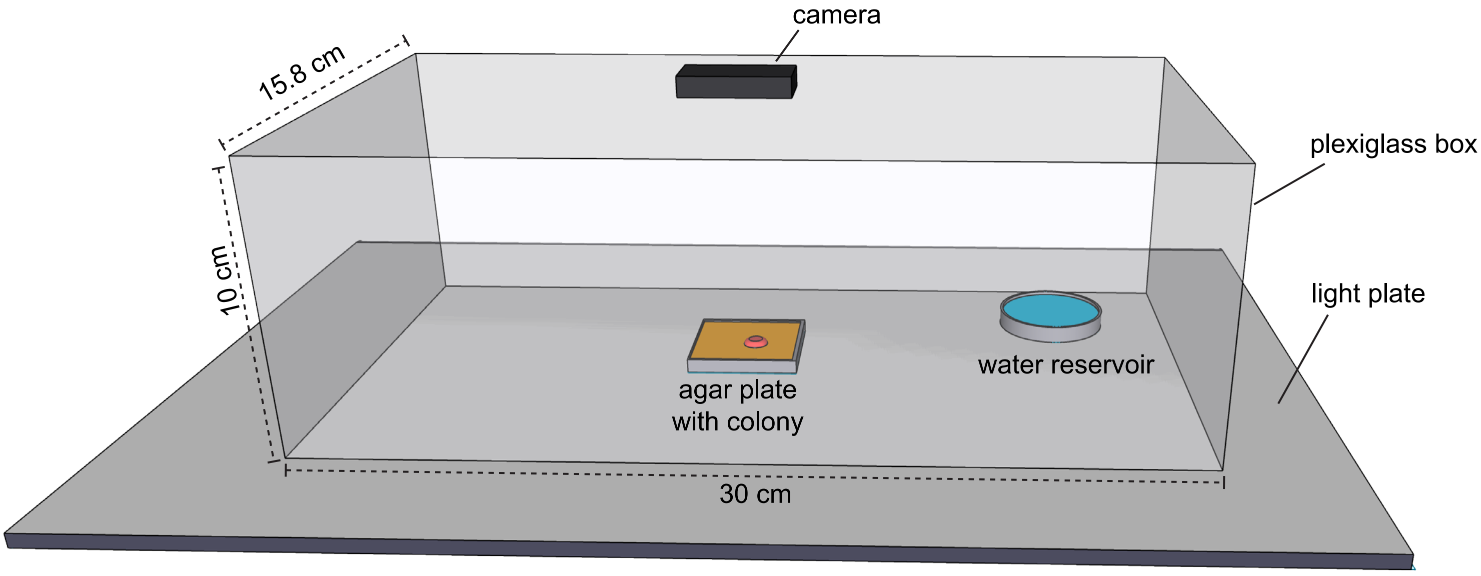


**Figure S1. (A)** Schematic showing the structure and reduction of TTC. **(B)** Schematic of incubation setup used to generate time-lapse movies of colony biofilm growth and TTC reduction.
