## Supplemental Figure 2 for "Light/dark and temperature cycling modulate metabolic electron flow in *Pseudomonas aeruginosa* biofilms"

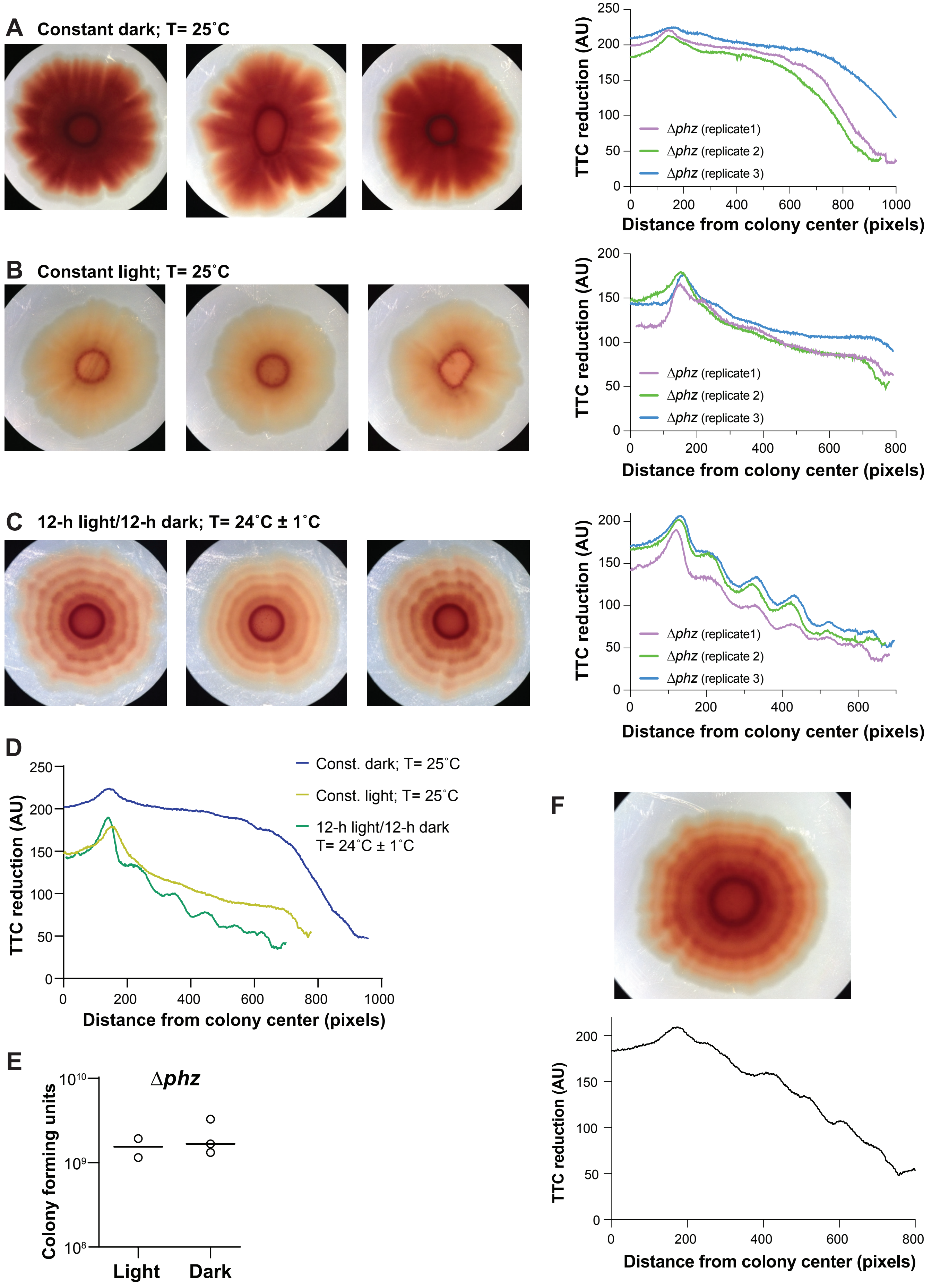

**Figure S2. (A-C)** Left: Replicates of Δ*phz* biofilms grown in constant dark at 25°C (A), in constant light at 25°C (B), and light and temperature cycling, 12-h light 25°C, 12-h dark 23°C (C). Right: Quantification of TTC reduction. **(D)** Comparison of representative TTC reduction quantification from all three conditions in panels (A-C). **(E)** Colony forming units for biofilms grown in constant light or constant dark at 25°C for 4 days. **(F)** Top: WT biofilm grown in light and temperature cycling, 12-h light 25 °C, 12-h dark 23°C . Bottom: Quantification of TTC reduction.
