## Supplemental Figure 3 for "Light/dark and temperature cycling modulate metabolic electron flow in *Pseudomonas aeruginosa* biofilms"

**Temperature cycling**

**Supplemental Figure 3**

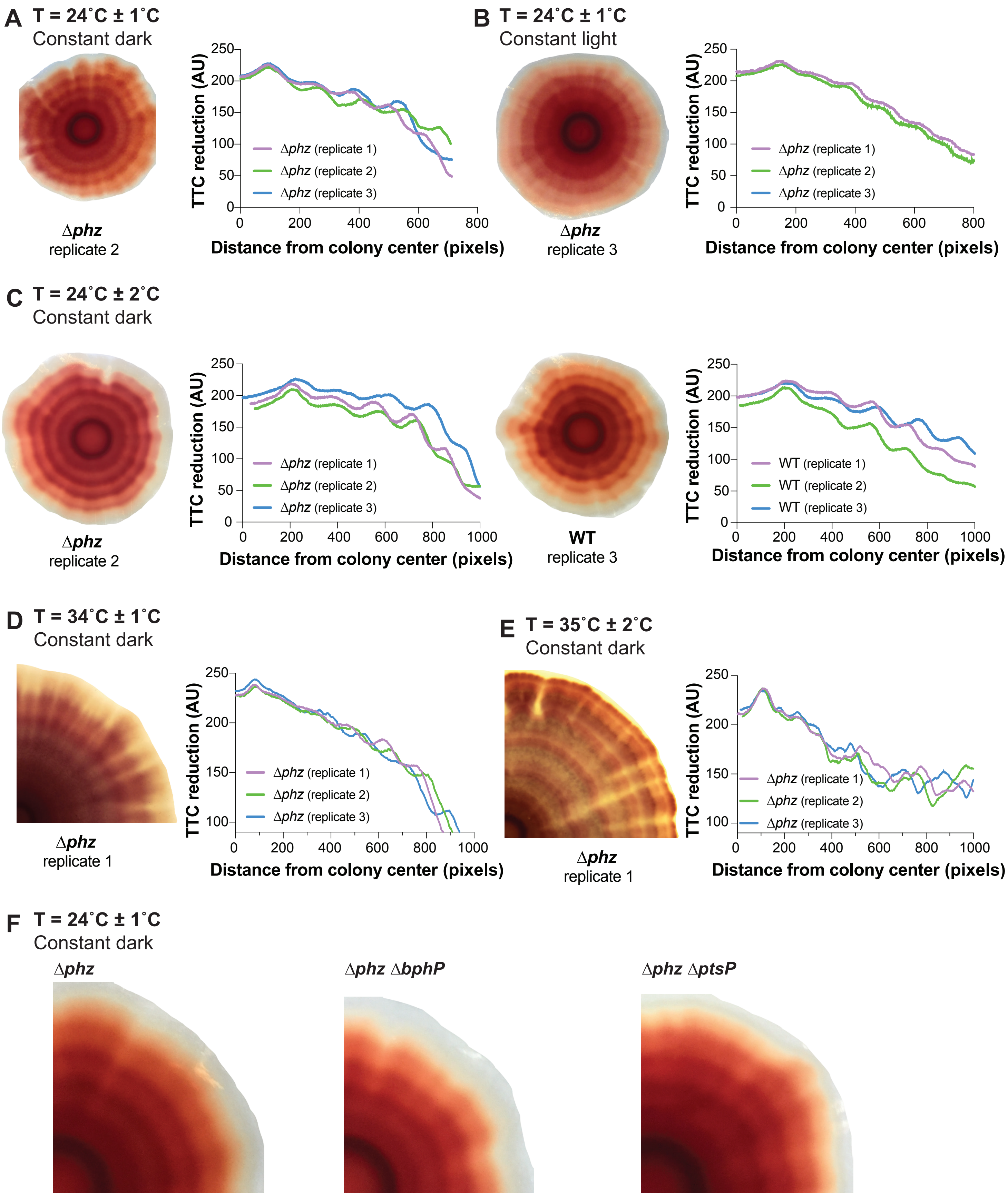

**Figure S3. (A)** Left: Representative replicate for Figure 1F. Right: Quantification of red color intensity (i.e., TTC reduction) at the indicated distance from the biofilm center for a radius of the biofilm for three replicates. **(B)** Left: Representative replicate for Δ*phz* biofilm grown in constant light with 24 °C +/- 1°C temperature cycling. Right: Quantification of red color intensity (i.e., TTC reduction) at the indicated distance from the biofilm center for a radius of the biofilm for three replicates. **(C)** Left: Representative WT or Δ*phz* biofilm grown in constant dark with 24°C +/- 2°C temperature cycling. Right: Quantification of TTC reduction for three replicates per condition. **(D&E)** Representative replicates for Δ*phz* biofilms grown in constant dark with 34°C +/- 1°C and 35°C +/- 2°C temperature cycling. **(F)** Light sensor mutants grown in constant dark with 24°C +/- 1°C temperature cycling, on medium containing 0.004% TTC.
