## Supplemental Figure 4 for "Light/dark and temperature cycling modulate metabolic electron flow in *Pseudomonas aeruginosa* biofilms"

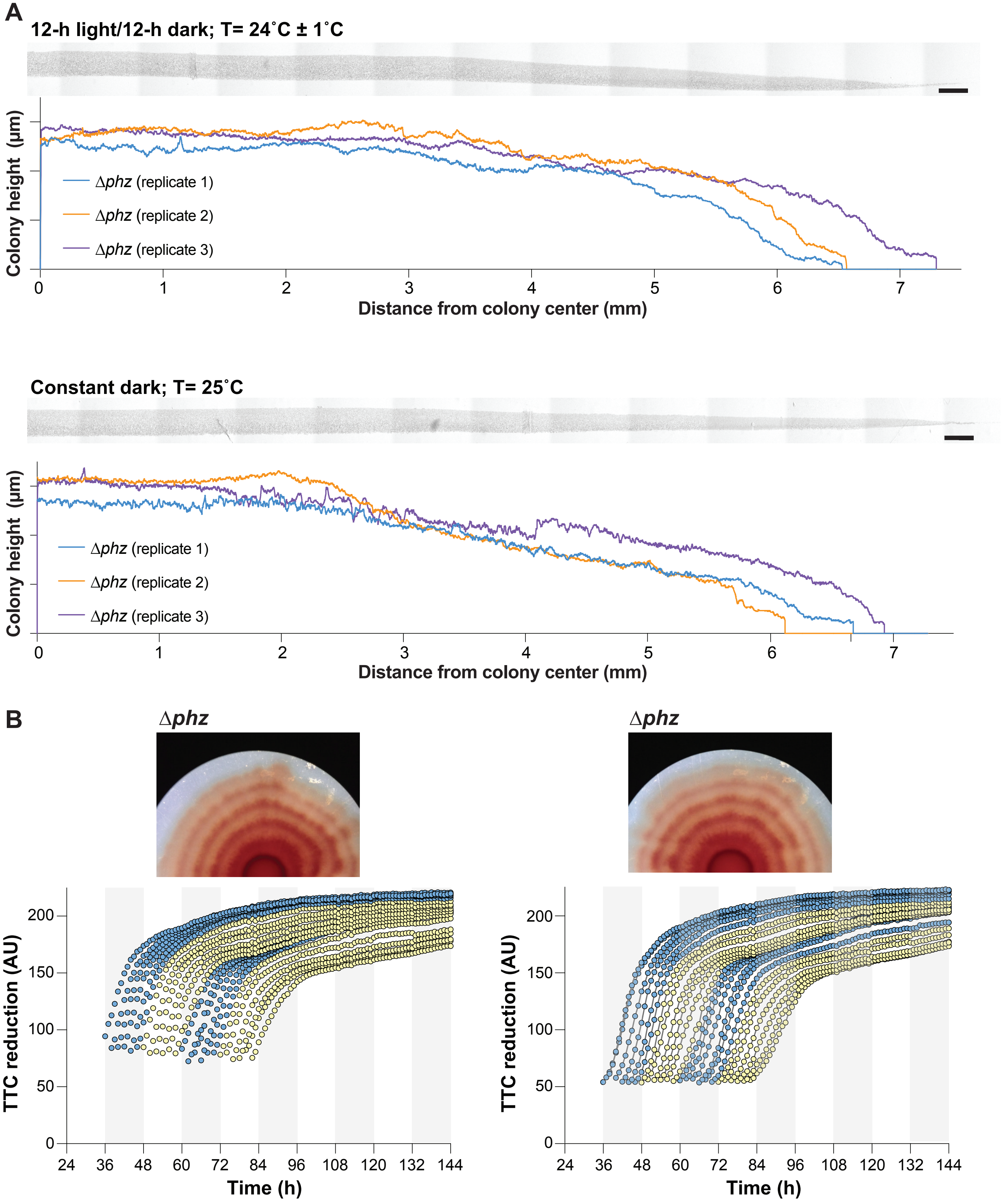

**Figure S4. (A)** Representative thin sections of Δ*phz* biofilms grown on medium containing 0.004% TTC for 3 days. Top: Section of biofilm grown with L/D and 24 ± 1 °C cycling. Bottom: Section of biofilm grown in the dark and at 25°C. Scale bars are 200 μm. For each condition, three replicates were quantified and graphed. Biological triplicates are shown. **(B)** Replicates for figure 2B. Traces corresponding to biomass appearing during dark intervals are shown in shades of blue; those corresponding to biomass appearing during light intervals are shown in shades of yellow.
