## Supplemental Figure 5 for "Light/dark and temperature cycling modulate metabolic electron flow in *Pseudomonas aeruginosa* biofilms"

**A** Principal component analysis

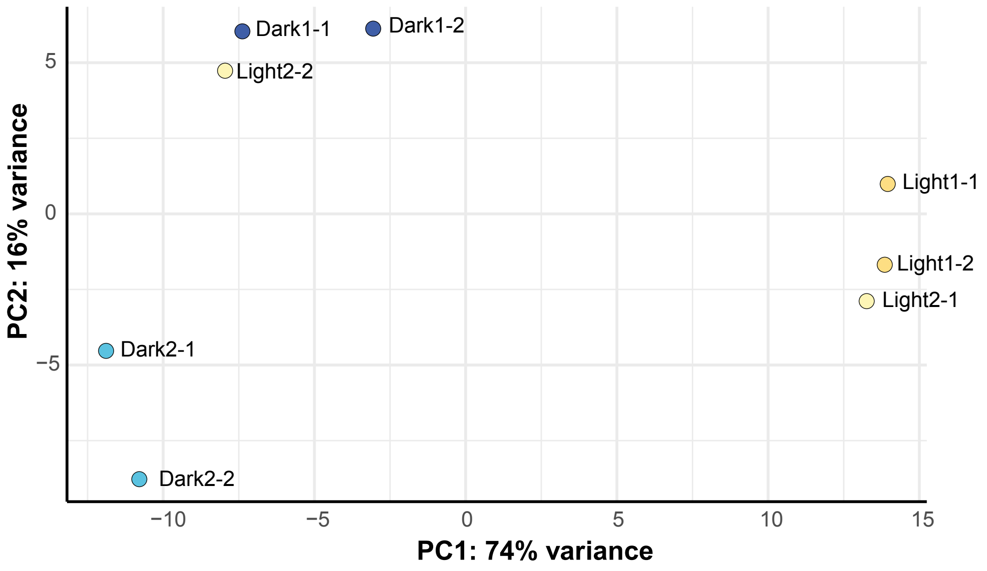


**B**

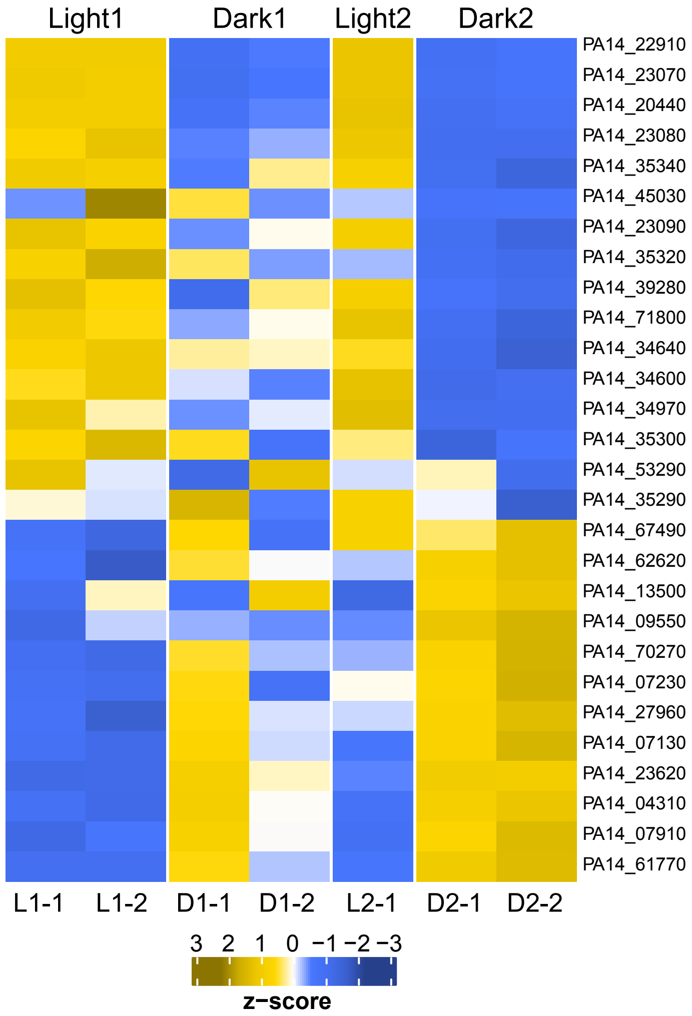


**C**

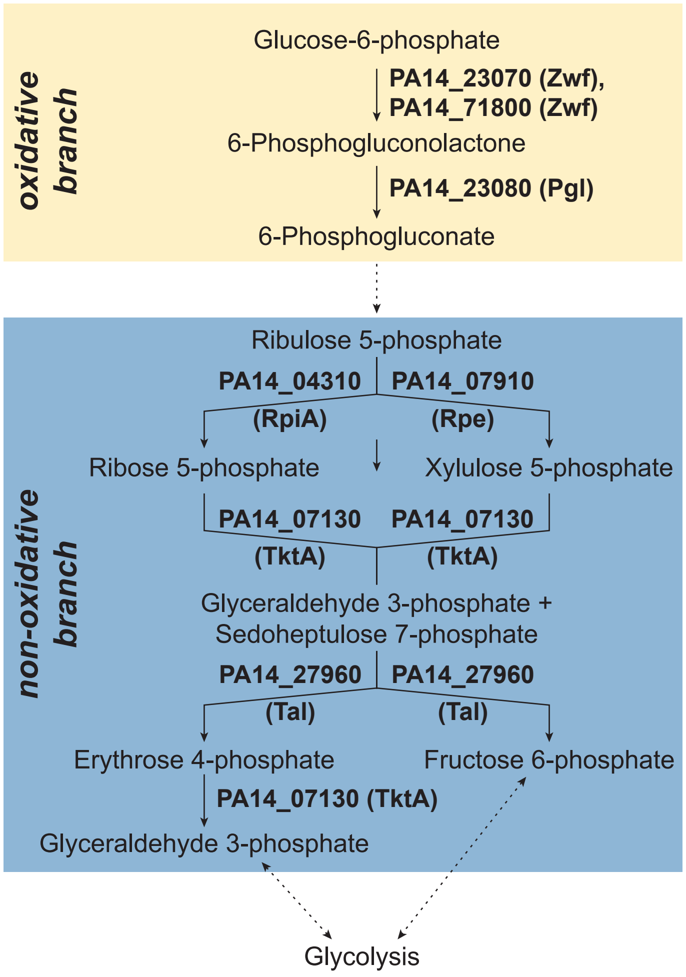

**Figure S5. (A)** Dimensionality reduction by principal component analysis was applied to RNAseq data for the full set of 8 samples obtained as shown in **Figure 6A**. Ninety percent of maximum possible variance was accounted for by the first two principle components PC1 and PC2. Each sample is plotted according to its values for PC1 and PC2. **(B)** Sixty-seven percent of genes involved in the pentose phosphate pathway were identified in the RNAseq as showing differential expression in biomass formed under light/dark and temperature cycling. Genes coding for enzymes involved in the oxidative branch were upregulated in biomass that formed under light/25 ˚C conditions, and genes coding for enzymes involved in the non-oxidative branch were upregulated in biomass that formed under dark/23˚C conditions. The heatmap shows genes involved in the pentose phosphate pathway, arranged according to their relative periodicity in samples from biomass formed under light/dark and temperature cycling. **(C)** Schematic of the pentose phosphate pathway highlighting selected genes whose expression was affected by growth under light/dark and temperature cycling conditions.
