## Supplemental Figure 6 for "Light/dark and temperature cycling modulate metabolic electron flow in *Pseudomonas aeruginosa* biofilms"

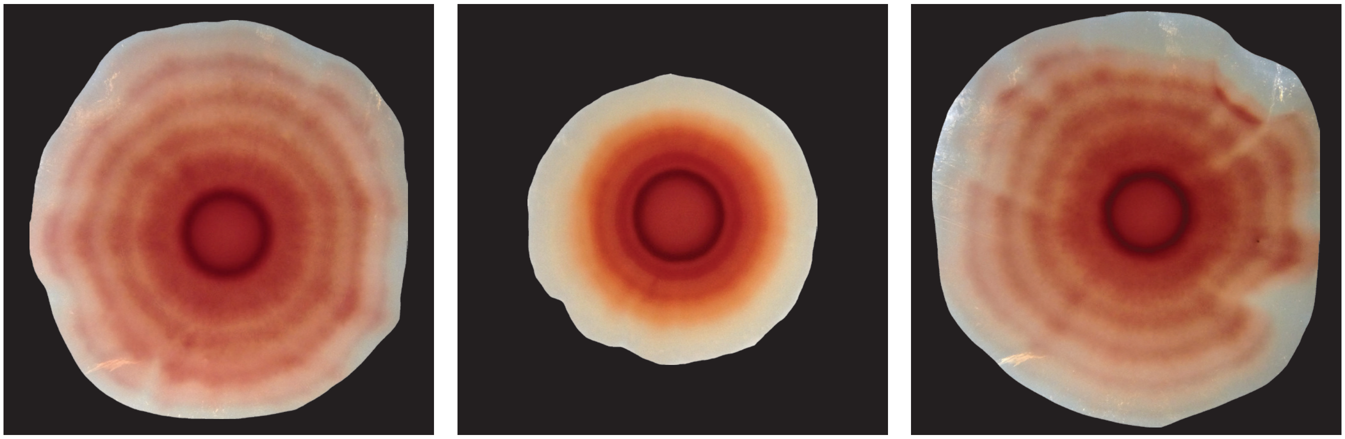


**Figure S6.** Images of Δ*phz*, Δ*phz*Δ*cco1cco2*, and Δ*phz* PaCco1 colony biofilms, corresponding to the analyses shown in Figure 7C and D.
