## Supplemental Table 1A-C and references for "Light/dark and temperature cycling modulate metabolic electron flow in *Pseudomonas aeruginosa* biofilms"

| Supplemental Table 1A: Strains used in This study | | | | |
|---|---|---|---|---|
| *Pseudomonas aeruginosa* strains | | | | |
| **Strain** | **Number** | **Description** | **Source** | **Used in figure** |
| PA14 Δ*phz* | LD24 | PA14 with deletions in *phzA1-G1* (*PA14_09480-09410*) and *phzA2-G2* (*PA14_39970-39880*) operons | (1) | 1C-G, 2B, 2C, 3, 4, 5, 6A-C(top) |
| PA14 Δ*phz* Δ*pel* | LD83 | PA14 Δ*phz* with deletions in genes *pelB-G* (*PA14_24490-24560*) | (2) | 3A, 3B |
| PA14 Δ*phz* Δ*bphP* | LD4480 | PA14 Δ*phz* with deletion in the bacteriophytochrome gene *bphP* (*PA14_10700*) | This study | 4 |
| PA14 Δ*phz* Δ*ptsP* | LD3131 | PA14 Δ*phz* with deletion in the phosphoenolpyruvate-protein transferase gene *ptsP* (*PA14_04410*) | This study | 4 |
| PA14 Δ*phz* Δ*cco1/2* | LD1938 | PA14 Δ*phz* with deletions of both *cco* operons (*PA14_44340-44400*) | (3) | 2D, 6C(mid) |
| PA14 Δ*phz* ΔPa*Cco1* | LD2104 | PA14 Δ*phz* with deletions in the *cox* operon (*PA14_01290-01320*), the *cyo* operon (*PA14_47150-47210*), the *cio* operon (*PA14_13030-13040*) and the *cco2* operon (*PA14_44340-44360*) | This study | 6C(bot) |
| *Escherichia coli* strains | | | | |
| UQ950 | LD44 | E. coli DH5 λ*pir* strain for cloning. F- Δ(*argF-lac*)169φ80 *dlacZ58*(Δ*M15*) *glnV44*(AS) *rfbD1 gyrA96*(Nal$^R$) *recA1 endA1 spoT thi-1 hsdR17 deoR* λ*pir*$^+$ | D. Lies | |
| BW29427 | LD661 | Donor strain for conjugation. thrB1004 pro thi rpsL hsdS lacZ ΔM15RP4-1360 Δ(araBAD)567 ΔdapA1314::[erm pir(wt)] | W. Metcalf, University of Illinois | |
| S17-1 | LD2901 | Str$^R$ , Tp$^R$ , F− RP4-2-Tc::Mu *aphA*::Tn7 *recA* λpir lysogen | (6) | |
| *Saccharomyces cerevisiae* strains | | | | |
| InvSc1 | LD676 | MATa/MATalpha leu2/leu2 trp1-289/ trp1-289 ura3-52/ ura3-52 his3-Δ1/his3-Δ1 | This study | |
| *Pseudomonas aeruginosa* strains used in screen for putative light sensors | | | | |
| **Strain** | **Number** | **Description** | **Source** | **Transposon plate location** |
| Δ*PA14_38740* | LD2594 | PA14 with deletion in the two-component sensor *PA14_38740* | This study | N/A |
| Δ*pilS* | LD2612 | PA14 with deletion in the two-component sensor gene *pilS* (*PA14_60250*) | This study | N/A |
| Δ*phz* Δ*prpR* | LD2593 | PA14 Δ*phz* with deletion in *prpR* (*PA14_36000*) | This study | N/A |

| | | | | |
|---|---|---|---|---|
| Δ*phz* Δ*pilS* | LD2613 | PA14 Δ*phz* with deletion in the two-component sensor gene *pilS* (*PA14_60250*) | This study | N/A |
| Δ*PA14_46860-46840* | LD696 | PA14 with deletion in *PA14_46860*, *PA14_46850*, and *PA14_46840* | This study | N/A |
| Δ*PA14_38970-38990* | LD2271 | PA14 with deletion in the two-component sensor gene *PA14_38970* and *PA14_38990* | This study | N/A |
| Δ*PA14_66320* | LD291 | PA14 with deletion in *PA14_66320* | (4) | N/A |
| Δ*phz* Δ*PA14_38740* | LD2595 | PA14 Δ*phz* with deletion in the two-component sensor gene *PA14_38740* | (4) | N/A |
| Δ*phz* Δ*PA14_66320* | LD290 | PA14 Δ*phz* with deletion in *PA14_66320* | (4) | N/A |
| Δ*PA14_40210* | LD2610 | PA14 with deletion in *PA14_40210* | This study | N/A |
| Δ*ntrB* | LD2594 | PA14 with deletion in *ntrB* (*PA14_67670*) | This study | N/A |
| Δ*phz* Δ*PA14_38970-38990* | LD2272 | PA14 Δ*phz* with deletion in the two-component sensor gene *PA14_38970* and *PA14_38990* | This study | N/A |
| ΔPA14_04420 | LD2077 | PA14 with deletion in *PA14_04420* | This study | N/A |
| Δ*PA14_46850* | LD694 | PA14 with deletion in the transcriptional regulator *PA14_04420* | This study | N/A |
| Δ*phz* Δ*PA14_40210* | LD2611 | PA14 Δ*phz* with deletion in *PA14_40210* | This study | N/A |
| Δ*rmcA* | LD2227 | PA14 with deletion in *rmcA* (*PA14_07500*) | (4) | N/A |
| Δ*PA14_48830* | LD1788 | PA14 with deletion in the transcriptional regulator *PA14_48830* | This study | N/A |
| Δ*phz* Δ*PA14_04420* | LD2078 | PA14 Δ*phz* with deletion in *PA14_04420* | This study | N/A |
| Δ*phz* Δ*PA14_48830* | LD1113 | PA14 Δ*phz* with deletion in the transcriptional regulator *PA14_48830* | This study | N/A |
| Δ*PA14_36000* | LD2592 | PA14 with deletion in *prpR* (*PA14_36000*) | This study | N/A |
| Δ*PA14_57170* | LD1338 | PA14 with deletions in the two component sensor gene *PA14_57170* | This study | N/A |
| Δ*phz* Δ*rmcA* | LD2228 | PA14 Δ*phz* with deletion in *rmcA* (*PA14_07500*) | (4) | N/A |
| Δ*phz* Δ*PA14_57170* | LD1339 | PA14 Δ*phz* with deletions in the two component sensor gene *PA14_57170* | This study | N/A |
| *PA14_02220::Tn* | | PA14 with a MAR2xT7 transposon insertion in the *PA14_02220* gene | (5) | PAMr_nr_mas_11_3, G7 |
| *PA14_10770::Tn* | | PA14 with a MAR2xT7 transposon insertion in the *PA14_10770* gene | (5) | PAMr_nr_mas_08_1, B5 |
| *PA14_21700::Tn* | | PA14 with a MAR2xT7 transposon insertion in the *PA14_21700* gene | (5) | PAMr_nr_mas_15_2, B12 |

| | | | | |
|---|---|---|---|---|
| *PA14_39560::Tn* | | PA14 with a MAR2xT7 transposon insertion in the *PA14_39560* gene | (5) | PAMr_nr_mas_6_3, B8 |
| *PA14_48160::Tn* | | PA14 with a MAR2xT7 transposon insertion in the *PA14_48160* gene | (5) | PAMr_nr_mas_9_1, H11 |
| *PA14_53140::Tn* | | PA14 with a MAR2xT7 transposon insertion in the *PA14_53140* gene | (5) | PAMr_nr_15_2, F3 |
| *PA14_59800::Tn* | | PA14 with a MAR2xT7 transposon insertion in the *PA14_59800* gene | (5) | PAMr_nr_mas_05_1, G11 |
| *PA14_65540::Tn* | LD1812 | PA14 with a MAR2xT7 transposon insertion in the *PA14_65540* gene | (5) | PAMr_nr_mas_03_2, C3 |
| *PA14_03720::Tn* | | PA14 with a MAR2xT7transposon insertion in the *PA14_03720* gene | (5) | PAMr_nr_mas_04_3, D7 |
| *PA14_10700::Tn* | | PA14 with a MAR2xT7 transposon insertion in the *PA14_10700* gene | (5) | PAMr_nr_mas_03_3, G2 |
| *PA14_24720::Tn* | | PA14 with a MAR2xT7 transposon insertion in the *PA14_24720* gene | (5) | PAMr_nr_mas_12_2, H12 |
| *PA14_39560::Tn* | | PA14 with a MAR2xT7 transposon insertion in the *PA14_39560* gene | (5) | PAMr_nr_mas_11_2, E10 |
| *PA14_49160::Tn* | | PA14 with a MAR2xT7 transposon insertion in the *PA14_49160* gene | (5) | PAMr_nr_mas_9_4, G9 |
| *PA14_53140::Tn* | | PA14 with a MAR2xT7 transposon insertion in the *PA14_53140* gene | (5) | PAMr_nr_mas_15_3, B4 |
| *PA14_59800::Tn* | | PA14 with a MAR2xT7 transposon insertion in the *PA14_59800* gene | (5) | PAMr_nr_mas_01_4, A5 |
| *PA14_65540::Tn* | LD1813 | PA14 with a MAR2xT7 transposon insertion in the *PA14_65540* gene | (5) | PAMr_nr_mas_07_4, B8 |
| *PA14_06950::Tn* | LD1444 | PA14 with a MAR2xT7 transposon insertion in the *PA14_06950* gene | (5) | ExMr_nr_mas_01_1, C10 |
| *PA14_11630::Tn* | | PA14 with a MAR2xT7 transposon insertion in the *PA14_11630* gene | (5) | PAMr_nr_mas_9_3, F9 |
| *PA14_32940::Tn* | | PA14 with a MAR2xT7 transposon insertion in the *PA14_32940* gene | (5) | PAMr_nr_mas_08_1, H7 |
| *PA14_44300::Tn* | | PA14 with a MAR2xT7 transposon insertion in the *PA14_44300* gene | (5) | PAMr_nr_mas_13_2, E11 |
| *PA14_50200::Tn* | LD715 | PA14 with a MAR2xT7 transposon insertion in the *PA14_50200* gene | (5) | PAMr_nr_mas_10_3, C5 |
| *PA14_53310::Tn* | | PA14 with a MAR2xT7 transposon insertion in the *PA14_53310* gene | (5) | PAMr_nr_mas_13_1, C3 |
| *PA14_60870::Tn* | | PA14 with a MAR2xT7 transposon insertion in the *PA14_60870* gene | (5) | PAMr_nr_mas_02_3, C10 |
| *PA14_70760::Tn* | | PA14 with a MAR2xT7 transposon insertion in the *PA14_70760* gene | (5) | PAMr_nr_mas_9_2, C1 |
| *PA14_07820::Tn* | | PA14 with a MAR2xT7 transposon insertion in the *PA14_07820* gene | (5) | PAMr_nr_mas_8_3, E1 |
| *PA14_11830::Tn* | | PA14 with a MAR2xT7 transposon insertion in the *PA14_11830* gene | (5) | PAMr_nr_mas_11_2, E4 |

| | | | | |
|---|---|---|---|---|
| *PA14_36420::Tn* | | PA14 with a MAR2xT7 transposon insertion in the *PA14_36420* gene | (5) | PAMr_nr_mas_9_1, B8 |
| *PA14_46030::Tn* | | PA14 with a MAR2xT7 transposon insertion in the *PA14_46030* gene | (5) | PAMr_nr_mas_8_2, E3 |
| *PA14_50200::Tn* | | PA14 with a MAR2xT7 transposon insertion in the *PA14_50200* gene | (5) | PAMr_nr_mas_12_4, F12 |
| *PA14_53310::Tn* | | PA14 with a MAR2xT7 transposon insertion in the *PA14_53310* gene | (5) | PAMr_nr_mas_13_2, A3 |
| *PA14_61640::Tn* | | PA14 with a MAR2xT7 transposon insertion in the *PA14_61640* gene | (5) | PAMr_nr_mas_06_2, H4 |
| *PA14_71850::Tn* | | PA14 with a MAR2xT7 transposon insertion in the *PA14_71850* gene | (5) | PAMr_nr_mas_04_3, D11 |
| *PA14_09680::Tn* | | PA14 with a MAR2xT7 transposon insertion in the *PA14_09680* gene | (5) | PAMr_nr_mas_11_3, E10 |
| *PA14_12820::Tn* | | PA14 with a MAR2xT7 transposon insertion in the *PA14_12820* gene | (5) | PAMr_nr_mas_9_4, F2 |
| *PA14_37690::Tn* | | PA14 with a MAR2xT7 transposon insertion in the *PA14_37690* gene | (5) | PAMr_nr_mas_14_1, B2 |
| *PA14_46850::Tn* | LD419 | PA14 with a MAR2xT7 transposon insertion in the *PA14_46850* gene | (5) | PAMr_nr_mas_04_3, E6 |
| *PA14_52980::Tn* | LD1445 | PA14 with a MAR2xT7 transposon insertion in the *PA14_52980* gene | (5) | PAMr_nr_mas_01_3, D6 |
| *PA14_55780::Tn* | | PA14 with a MAR2xT7 transposon insertion in the *PA14_55780* gene | (5) | PAMr_nr_mas_11_4, B9 |
| *PA14_62530::Tn* | LD411 | PA14 with a MAR2xT7 transposon insertion in the *PA14_62530* gene | (5) | PAMr_nr_mas_06_2, A7 |
| *PA14_10290::Tn* | | PA14 with a MAR2xT7 transposon insertion in the *PA14_10290* gene | (5) | PAMr_nr_mas_05_4, D5 |
| *PA14_21700::Tn* | | PA14 with a MAR2xT7 transposon insertion in the *PA14_21700* gene | (5) | PAMr_nr_mas_07_2, C4 |
| *PA14_38570::Tn* | | PA14 with a MAR2xT7 transposon insertion in the *PA14_38570* gene | (5) | PAMr_nr_mas_11_2, H11 |
| *PA14_47910::Tn* | | PA14 with a MAR2xT7 transposon insertion in the *PA14_47910* gene | (5) | PAMr_nr_mas_03_4, E7 |
| *PA14_53140::Tn* | | PA14 with a MAR2xT7 transposon insertion in the *PA14_53140* gene | (5) | PAMr_nr_mas_10_4, B7 |
| *PA14_59780::Tn* | | PA14 with a MAR2xT7 transposon insertion in the *PA14_59780* gene | (5) | PAMr_nr_mas_15_1, G10 |
| *PA14_62530::Tn* | LD412 | PA14 with a MAR2xT7 transposon insertion in the *PA14_62530* gene | (5) | PAMr_nr_mas_15_1, F6 |
| *PA14_72390::Tn* | | PA14 with a MAR2xT7 transposon insertion in the *PA14_72390* gene | (5) | PAMr_nr_mas_07_1, D4 |
| *PA14_04410::Tn* | | PA14 with a MAR2xT7 transposon insertion in the *PA14_04410* gene | (5) | PAMr_nr_mas_14_3, A11 |
| *PA14_29620::Tn* | | PA14 with a MAR2xT7 transposon insertion in the *PA14_29620* gene | (5) | PAMr_nr_mas_01_2, E8 |

| | | | | |
|---|---|---|---|---|
| *PA14_43350::Tn* | | PA14 with a MAR2xT7 transposon insertion in the *PA14_43350* gene | (5) | PAMr_nr_mas _05_2, A3 |
| *PA14_10190::Tn* | | PA14 with a MAR2xT7 transposon insertion in the *PA14_10190* gene | (5) | PAMr_nr_mas _08_4, F10 |
| *PA14_29620::Tn* | | PA14 with a MAR2xT7 transposon insertion in the *PA14_29620* gene | (5) | PAMr_nr_mas _10_4, G2 |
| *PA14_43350::Tn* | | PA14 with a MAR2xT7 transposon insertion in the *PA14_43350* gene | (5) | PAMr_nr_mas _12_4, H3 |
| *PA14_23190::Tn* | | PA14 with a MAR2xT7 transposon insertion in the *PA14_23190* gene | (5) | PAMr_nr_mas _11_4, G7 |
| *PA14_31330::Tn* | | PA14 with a MAR2xT7 transposon insertion in the *PA14_31330* gene | (5) | PAMr_nr_mas _03_2, B11 |
| *PA14_43430::Tn* | | PA14 with a MAR2xT7 transposon insertion in the *PA14_43430* gene | (5) | PAMr_nr_mas _09_3, D10 |
| *PA14_02910::Tn* | | PA14 with a MAR2xT7 transposon insertion in the *PA14_02910* gene | (5) | PAMr_nr_mas _05_2, A4 |
| *PA14_24510::Tn* | | PA14 with a MAR2xT7 transposon insertion in the *PA14_24510* gene | (5) | PAMr_nr_mas _09_4, B3 |
| *PA14_38500::Tn* | | PA14 with a MAR2xT7 transposon insertion in the *PA14_38500* gene | (5) | PAMr_nr_mas _12_3, B7 |
| *PA14_56430::Tn* | | PA14 with a MAR2xT7 transposon insertion in the *PA14_56430* gene | (5) | PAMr_nr_mas _13_3, E7 |
| *PA14_04410::Tn* | | PA14 with a MAR2xT7 transposon insertion in the *PA14_04410* gene | (5) | PAMr_nr_mas _05_2, G7 |
| *PA14_27570::Tn* | | PA14 with a MAR2xT7 transposon insertion in the *PA14_27570* gene | (5) | PAMr_nr_mas _02_1, C11 |
| *PA14_38500::Tn* | | PA14 with a MAR2xT7 transposon insertion in the *PA14_38500* gene | (5) | PAMr_nr_mas _15_3, E10 |
| *PA14_04410::Tn* | | PA14 with a MAR2xT7 transposon insertion in the *PA14_04410* gene | (5) | PAMr_nr_mas _07_2, G7 |
| *PA14_28070::Tn* | | PA14 with a MAR2xT7 transposon insertion in the *PA14_28070* gene | (5) | PAMr_nr_mas _06_4, F7 |
| *PA14_42970::Tn* | | PA14 with a MAR2xT7 transposon insertion in the *PA14_42970* gene | (5) | PAMr_nr_mas _04_3, H5 |

**Supplemental Table 1B: Primers used in This study**

| **Primer number** | **Sequence** | **used to make plasmid number** |
|---|---|---|
| LD2475 | aggcaaattctgttttatcagaccgcttctgcgttctgatCGGCTCGATCACTTCCTGCA | pLD3125 |
| LD2476 | ctgcgggttgtcgaaggtgagCATGGCTTCCTTGACCCGCTG | |
| LD2477 | cagcgggtcaaggaagccatgCTCACCTTCGACAACCCGCAG | |

| | | |
|---|---|---|
| LD2478 | ggaattgtgagcggataacaatttcacacaggaaacagctGATGAACTCCTCGCCGCCC | |
| LD3030 | ggaattgtgagcggataacaatttcacacaggaaacagctGCGCGGGGATGCCCATTTAT | pLD3618 |
| LD3031 | gttgcggttgcgctggttgtagttcgccagggtaaccgggg | |
| LD3032 | ccccggttaccctggcgaactacaaccagcgcaaccgcaac | |
| LD3033 | caaattctgttttatcagaccgcttctgcgttctgatCTCGTTCCGCGGAGTCGCCG | |
| LD1833 | ccaggcaaattctgttttatcagaccgcttctgcgttctgatGGCGCGGTACTTTCACTC | pLD2622 |
| LD1834 | tgtccagcgtctcctgtatgtGGACTCCCATGGCTTCCT | |
| LD1835 | aggaagccatgggagtccACATACAGGAGACGCTGGACA | |
| LD1836 | ggaattgtgagcggataacaatttcacacaggaaacagctCTGCGGCTGACCCTCAGT | |
| LD1845 | ccaggcaaattctgttttatcagaccgcttctgcgttctgatGAGTTCGCCCAGCTCACC | pLD2604 |
| LD1846 | ctggttgctctcgcagagttGGACCTGATCTTCCAGTTCG | |
| LD1847 | cgaactggaagatcaggtccAACTCTGCGAGAGCAACCAG | |
| LD1848 | ggaattgtgagcggataacaatttcacacaggaaacagctTTCTTCAGCTTCTCCTGTGC | |
| LD39 | GGAATTGTGAGCGGATAACAATTTCACACAGGAAACAGCTGGTCGC GGATATAACCTGAA | pLD4558 |
| LD40 | CAGGTAGTCGATCAGTGCCGGACAAAGCTCGGAAAGACGA | |
| LD41 | TCGTCTTTCCGAGCTTTGTCCGGCACTGATCGACTACCTG | |
| LD42 | CCAGGCAAATTCTGTTTTATCAGACCGCTTCTGCGTTCTGACTACGA CATGGCGATCCTG | |
| LD1344 | ccaggcaaattctgttttatcagaccgcttctgcgttctgatGACTTCGCCGCCTACCTG | pLD2267 |
| LD1345 | gaagtggtggcccaggacCGAGCTTGGCCATACCCTGAT | |
| LD1346 | atcagggtatggccaagctcgGTCCTGGGCCACCACTTC | |
| LD1347 | ggaattgtgagcggataacaatttcacacaggaaacagctGCTCGAACATCTTCCTCGAC | |
| LD1841 | ccaggcaaattctgttttatcagaccgcttctgcgttctgatCTACCGGCCGATGTACTACC | pLD2624 |
| LD1842 | agagggtcatgcgggtccCAGGGACGAAGAAGGGAGAG | |
| LD1843 | ctctcccttcttcgtccctgGGACCCGCATGACCCTCT | |
| LD1844 | ggaattgtgagcggataacaatttcacacaggaaacagctGAAGATCACCGCCTACGACT | |
| LD1849 | ccaggcaaattctgttttatcagaccgcttctgcgttctgatCTGGACGGCAAACCCTAC | pLD2605 |
| LD1850 | gctggctgatgatgttctggATGTACTCCAGGCGCAGTTC | |
| LD1851 | gaactgcgcctggagtacatCCAGAACATCATCAGCCAGC | |
| LD1852 | ggaattgtgagcggataacaatttcacacaggaaacagctCTTCAGCAGCTTGGGTTCC | |
| LD1059 | GGAATTGTGAGCGGATAACAATTTCACACAGGAAACAGCTGCGAGC TGAGCAAGGGCCTG | pLD2079 |
| LD1060 | TCAGCAACAGGCCACGCAATGTAAGCGCCGCACGACGAAG | |
| LD1062 | CTTCGTCGTGCGGCGCTTACATTGCGTGGCCTGTTGCTGA | |

| | | |
|---|---|---|
| LD1061 | CCAGGCAAATTCTGTTTTATCAGACCGCTTCTGCGTTCTGATGACGATGATCAGGCCGCCGC | |
| LD306 | ggaattgtgagcggataacaatttcacacaggaaacagctCAGCAGCAGAACGAAACTCA | pLD1328 |
| LD307 | CTGGTTGGTGGACTGCCTGCGCAACCTGATAGAAGACG | |
| LD308 | CGTCTTCTATCAGGTTGCGCAGGCAGTCCACCAACCAG | |
| LD309 | ccaggcaaattctgttttatcagaccgcttctgcgttctgatCATGGTTCAGAAGCGCAGT | |

#### Supplemental Table 1C: Plasmids used in This study

| Number | Description | Source |
|---|---|---|
| pMQ30 | 7.5 kb mobilizable vector; oriT, sacB, GmR | (7) |
| pLD3125 | Δ*ptsP* (Δ*PA14_04410*) PCR fragment introduced into pMQ30 by gap repair cloning in yeast strain InvSc1 | This study |
| pLD3618 | Δ*bphP* (Δ*PA14_10700*) PCR fragment introduced into pMQ30 by gap repair cloning in yeast strain InvSc1 | This study |
| pLD2623 | Δ*PA14_38740* (Δ*PA14_38740*) PCR fragment introduced into pMQ30 by gap repair cloning in yeast strain InvSc1 | This study |
| pLD2604 | Δ*PA14_60250* (Δ*PA14_60250*) PCR fragment introduced into pMQ30 by gap repair cloning in yeast strain InvSc1 | This study |
| pLD2622 | Δ*PA14_36000* (Δ*PA14_36000*) PCR fragment introduced into pMQ30 by gap repair cloning in yeast strain InvSc1 | This study |
| pLD2627 | Δ*PA14_38970-38990* (Δ*PA14_38970-38990*) PCR fragment introduced into pMQ30 by gap repair cloning in yeast strain InvSc1 | This study |
| pLD216 | Δ*PA14_66320* (Δ*PA14_66320*) PCR fragment introduced into pMQ30 by gap repair cloning in yeast strain InvSc1 | (4) |
| pLD2623 | Δ*PA14_38740* (Δ*PA14_38740*) PCR fragment introduced into pMQ30 by gap repair cloning in yeast strain InvSc1 | (4) |
| pLD1308 | Δ*PA14_03790* (Δ*PA14_03790*) PCR fragment introduced into pMQ30 by gap repair cloning in yeast strain InvSc1 | (4) |
| pLD2624 | Δ*PA14_40210* (Δ*PA14_40210*) PCR fragment introduced into pMQ30 by gap repair cloning in yeast strain InvSc1 | This study |
| pLD2605 | Δ*PA14_67670* (Δ*PA14_67670*) PCR fragment introduced into pMQ30 by gap repair cloning in yeast strain InvSc1 | This study |
| pLD2079 | Δ*PA14_04420* (Δ*PA14_04420*) PCR fragment introduced into pMQ30 by gap repair cloning in yeast strain InvSc1 | This study |
| pLD909 | Δ*rmcA* (Δ*PA14_07500*) PCR fragment introduced into pMQ30 by gap repair cloning in yeast strain InvSc1 | (4) |
| pLD2079 | Δ*PA14_04420* (Δ*PA14_04420*) PCR fragment introduced into pMQ30 by gap repair cloning in yeast strain InvSc1 | (4) |

### SUPPLEMENTAL MATERIAL REFERENCES
