## Supplemental file 2 for "Light/dark and temperature cycling modulate metabolic electron flow in *Pseudomonas aeruginosa* biofilms"

#!/usr/bin/env python
# coding: utf-8

# In[146]:


import numpy as np
import pandas as pd
import os
import math
from numba import jit
from PIL import Image
from scipy.optimize import curve_fit
import matplotlib.pyplot as plt


# In[147]:


#We want to write a function where agar = 0 and colony = 1, given a certain thershold. Make a new image. 
def colony_image(image_as_array, threshold):
    #Let's make a blank matrix to put the image in. 
    new_image = np.zeros(np.shape(image_as_array))
    
    #We want to loop through the image and check if values are above or below the threshold. 
    for row in range(len(image_as_array)):
        for column in range(len(image_as_array[0])):
            if image_as_array[row][column] < threshold:
                new_image[row][column] = 1 #below threshold is part of the colony. 
            else:
                new_image[row][column] = 0 
    return new_image


# In[148]:


@jit 
def find_left_border(col_image):
    shape = np.shape(col_image)
    left_border = np.zeros(shape[0])
    
    for row in range(len(col_image)):
        for column in range(len(col_image[0])):
            if col_image[row][column] == 0:
                try:
                    if np.mean(col_image[row][column:column+20]) == 0.95: #start at first entry and end at the entry before your last ie entry 0-19
                        left_border[row] = column
                except:
                    pass
    return left_border


# In[149]:


@jit 
def find_right_border(col_image):
    shape = np.shape(col_image)
    right_border = np.zeros(shape[0])
    
    for row in range(len(col_image)):
        for column in range(len(col_image[0])):
            if col_image[row][column] == 1:
                try:
                    if np.mean(col_image[row][column:column+20]) == 0.95: #start at first entry and end at the entry before your last ie entry 0-19
                        if col_image[row][column+19] == 0:
                            right_border[row] = column + 19
                except:
                    pass
    return right_border


# In[150]:


def find_left_transitions(image_as_array, threshold):
    ci = colony_image(image_as_array, threshold)
    left_transitions = find_left_border(ci)
    return left_transitions


# In[151]:


def find_right_transitions(image_as_array, threshold):
    ci = colony_image(image_as_array, threshold)
    right_transitions = find_right_border(ci)
    return right_transitions


# In[152]:


def refine_threshold (image_as_array, initial_guess):
    #We do this by estimating the borders this an initial guess. 
    #Evaluating the points halfway to the left as agar to get agar stats and refining the border from there.  
    left_border = find_left_transitions(image_as_array, initial_guess)
    right_border = find_right_transitions(image_as_array, initial_guess)
    #halfway to left border is the pixel value cut in half
    half_left = left_border/2
    #right is more complicated, we need the distance from the edge to the border cut in two and added back to the border position. 
    right_edge = len(image_as_array[0]) - 1
    distance = right_edge - right_border
    half_right = right_border + distance/2
    #Get stats of agar
    agar_values = np.empty(0) #empty array to store agar values in. 
    for row in range(len(image_as_array)):
        if left_border[row] >0 and right_border[row] >0: 
            for value_index in range(len(image_as_array[0])):
                if value_index <= half_left[row] or value_index >= half_right[row]:
                    value = image_as_array[row][value_index]
                    agar_values = np.append(agar_values, value)
    #calculate agar value stat
    agar_mean = np.mean(agar_values)
    agar_std = np.std(agar_values)
    refined_threshold = np.floor(agar_mean - 3*agar_std)
    return refined_threshold


# In[153]:


@jit (nopython=True)
def RGB_to_gray(im):
    shape = np.shape(im)
    gray_image = np.zeros((shape[0], shape[1]))
    for row in range(shape[0]):
        for column in range(shape[1]):
            red = im[row][column][0]
            green = im[row][column][1]
            blue = im[row][column][2]
            gray = int(.299*red + .587*green+.114*blue)
            #gray = int(0.5*red + 0.25*green+ 0.25*blue)
            if gray > 255:
                gray = 255
            gray_image[row][column] = gray
    return gray_image


# In[154]:


#First step is generate a stack (of images). 
#Tell python what folder to look in and then dump the image files. 
#Directory is just a string so I can cancatinate the names to get to the folder I want. 
def generate_stack(folder_name): #this will generate a stack with all the images in a sub-directory.
    parent_folder = os.getcwd()
    directory = parent_folder + "\\" + folder_name #this is windows, on mac/linux need +"/" instead
    stack = [] #This is the stack object that we want in the end. 
    
    #\\ Generate a list of file names
    
    filenames = [] #create a list of file names. 
    for file in os.listdir(directory):
        if file.endswith(".tif"):
            filenames.append(file) #here the files are not in order
    filenames.sort() #this will sort files - tried alpha then numerically but they only change in number so it is ohkay.
    #\\
    
    #\\ the next thing to do is load each file one by one and turn it into a matrix. Then store the matrix in stack.
    #load images. 
    for image in range(len(filenames)):
        name = directory + "\\" + filenames[image]
        img = Image.open(name, mode ="r") #open one image and convert to greyscale
        matrix = np.asarray(img) #turn image into matrix
        matrix = RGB_to_gray(matrix)
        stack.append(matrix)
    stack2 = np.asarray(stack) #turn the stack from a list to an array. 
    return stack2


# In[155]:


def generate_time_matrix(stack, initial_guess): #return an n by m matrix where value corresponds to slice number where pixel was 
    #first detected in the colony. Guess is for the stats. 
    shape = np.shape(stack)
    last_slice = stack[shape[0]-1]
    threshold = refine_threshold_n(last_slice, initial_guess)
    
    #now we have a thershold to analyze the borders of every image with. 
    tm = np.full((shape[1], shape[2]), -1) #shape 1 = number of rows, shape 2 = number of coulmns. Made a matrix the same size
    #as every image and filled it with -1's. 
    
    #\\ loop thorugh every image. 
    for index in range(shape[0]): #index for every slice. 
        #once we are in a image we want it's borders. 
        current_image = stack[index]
        border_left = find_left_transitions(current_image, threshold)
        border_right = find_right_transitions(current_image, threshold)
        
        #If a pixel is inbetweent he two borders, it is part of the colony. 
        #If we find no border then the border is = o from our transition function, so need to ignore those values. 
        #Because the image has mulitple rows, ther eis a value in each border for every row - need to loop through rows. 
        for row in range(len(border_left)):
            if border_left[row] > 0 and border_right[row] > 0:
                location1 = int(border_left[row])
                location2 = int(border_right[row]) #we are interested in the columns from location1 -> location2. 
                for column in range(location1, location2):
                    #we are only interested in, if this value has been stored in our time matrix, but the only way to have a 
                    #value of -1 (all slices are positive) is if nothing was stored. 
                    if tm[row][column] == -1: #if nothing was stored before...
                        tm[row][column] = index #index is slice number and therefore time. 
    print(threshold)
    return tm               


# In[156]:


def get_area (array):
    area_array = np.empty(0) #made emptry array
    for numbers in range(1, len(array)+1):
        area = np.sum(array[0:numbers])# : does not include the second position so have to start one later and end one later. 
        area_array = np.append(area_array, area) #append areas from each loop to empty array  with np.append(where to appeand, what to appead)
    return area_array


# In[157]:


def get_radius (area):
    pi = math.pi
    radius_array = np.empty(0) #made an empty array
    for index in range(len(area)):
        #at area one want to do area rad(one/pi)
        radius = (area[index]/pi)**(1/2) #area[index] means the value of area at index x (ie area[2] is value of are at index 2 so position 3).
        radius_array = np.append(radius_array, radius)
    return radius_array


# In[158]:


def colony_statistics(new_pixels_array, first_time_point, time_interval): #return new pixels for each timepoint, area, and radius (assume biofilm is a circle)
    length_of_movie = len(new_pixels_array)
    #We need to generate a time array
    time_points = np.zeros(length_of_movie)
    for time in range(length_of_movie):
        time_points[time] = first_time_point + time_interval*time
    area = get_area(new_pixels_array)
    radius = get_radius(area)
    
    #Now we need ot put together everything into a dataframe.
    dataframe = pd.DataFrame({"Time": time_points, "new_pixels" : new_pixels_array, "area" : area, "radius" : radius})
    return dataframe 


# In[159]:


def highlight_border(image_as_array, threshold):
    transitions_left = find_left_transitions(image_as_array, threshold)
    transitions_right = find_right_transitions(image_as_array, threshold)
    edited_image = image_as_array.copy("C")
    #want to loop through every row and turn the transition pixel black
    for row in range(len(image_as_array)): #loop through every row
        left_pixel = int(transitions_left[row]) 
        left_pixel_2 = int(transitions_left[row] -1)
        right_pixel = int(transitions_right[row])
        right_pixel_2 = int(transitions_right[row] +1)
        edited_image[row][left_pixel] = 0
        edited_image[row][left_pixel_2] = 0
        edited_image[row][right_pixel] = 0
        edited_image[row][right_pixel_2] = 0 
    return edited_image


# In[160]:


def filter_movie(image_stack, x_images): #example if x_images = 2 you take every other image. 
    running_stack = []
    for image_index in range(0, len(image_stack), x_images): #still makes an index, but giving range() more information. 
        image = image_stack[image_index]
        running_stack.append(image)
    #want to turn running_stack into an array
    array = np.asarray(running_stack)
    return array


# In[161]:


#time matrix, image_stack(movie), and how many hours forward to look. This is because a pixel location in the time matrix, 
#is in the same location as in a movie image. Will end up needing to truncate because if they haven't assymptoted they look like
#they reduced more TTC.  
def generate_reduction_matrix(image_stack, time_matrix, images_forward, initial_guess):
    #Returned an n by m matrix where the i, j entry is the percent TTC reduced by pixel i, j after images_forward amount of 
    #time compared to the end of the movie. 
    
    #We wanted to loop through the time matrix, looking at any pixel whose value >-1 ( use -1 as place holder for not colony)
    #And then the pixel vlaue x images_forward and compare it to the difference bwtn the threshold (starting iamge) 
    #and the endpoint value. 
    
    shape = np.shape(image_stack)
    last_image = image_stack[shape[0]-1]
    threshold = refine_threshold_n(last_image, initial_guess)
    
    #loop through the time matrix
    shape2 = np.shape(time_matrix)
    reduction_matrix = np.zeros(shape2) #empty matrix to populate. 
    point_max = shape[0] - images_forward
    
    for row in range(len(time_matrix)):
        for column in range(len(time_matrix[0])):
            point = time_matrix[row][column]
            if point >= 0 and point < point_max:
                points_forward = np.int(point+images_forward)
                ending_value = last_image[row][column]
                difference = threshold - ending_value 
                value_here = image_stack[points_forward][row][column]
                difference_here = threshold - value_here
                percentage = difference_here/difference *100 
                reduction_matrix[row][column] = percentage
                
            else:
                reduction_matrix[row][column] = -1 #-1 is placeholder. 
    return reduction_matrix


# In[162]:


#have a reduction matrix, gives a 2-D image. If we want to compile it into a 1-D image to make, say a graph
#we need to average the pixels which appear in a given time. 
#will split across two functions. 
@jit(nopython=True)
def generate_list_of_lists(time_matrix, reduction_matrix, image_stack):
    bigger_list = []
    
    #we want to loop through the time matrix for every time point 
    #For each time_point we want to generate a list of all perentages at that time
    #we want ot appeand that smaller list to our bigger list. precentage
    
    shape = np.shape(image_stack)
    
    for time in range(shape[0]): #loop through time points
        small_list = []
        for row in range(len(time_matrix)):
            for column in range(len(time_matrix[0])):
                if time_matrix[row][column] == time:
                    small_list.append(reduction_matrix[row][column])
        bigger_list.append(small_list)
    return bigger_list


# In[163]:


def parse_list_of_lists (list_of_lists):
    
    #We want the average and standard deviation 
    #We loop through the list of lists and cacluate the avg and std dev for each image (hours)
    times = len(list_of_lists) #Number of images. 
    
    #empty arrays to fill in values. 
    time_points = np.zeros(times)
    average = np.zeros(times)
    standard_deviation = np.zeros(times)
    
    for index in (range(times)): # looop through our list of lists. 
        time_points[index] = index 
        avg = np.mean(list_of_lists[index])
        std = np.std(list_of_lists[index])
        average [index] = avg
        standard_deviation[index] = std
    
    data = pd.DataFrame({"Time (hr) pixel entered colony": time_points, "Average % TTC Reduced" :average , "Standard Error" : standard_deviation})
    return data


# In[164]:


#time matrix, image_stack(movie), and how many hours forward to look. This is because a pixel location in the time matrix, 
#is in the same location as in a movie image. Will end up needing to truncate because if they haven't assymptoted they look like
#they reduced more TTC.  
def generate_TTC_here_matrix(image_stack, time_matrix, images_forward, initial_guess):
    #Returned an n by m matrix wher the i, j entry is the percent TTC reduced by pixel i, j after images_forward amount of 
    #time compared to the end of the movie. 
    
    #We wanted to loop through the time matrix, looking at any pixel whose value >-1 ( use -1 as place holder for not colony)
    #And then the pixel vlaue x images_forward and compare it to the difference bwtn the threshold (starting iamge) 
    #and the endpoint value. 
    
    shape = np.shape(image_stack)
    last_image = image_stack[shape[0]-1]
    threshold = refine_threshold(last_image, initial_guess)
    
    #loop through the time matrix
    shape2 = np.shape(time_matrix)
    reduction_matrix = np.zeros(shape2) #empty matrix to populate. 
    point_max = shape[0] - images_forward
    
    for row in range(len(time_matrix)):
        for column in range(len(time_matrix[0])):
            point = time_matrix[row][column]
            if point >= 0 and point < point_max:
                points_forward = np.int(point+images_forward)
                ending_value = last_image[row][column]
                difference = threshold - ending_value 
                value_here = image_stack[points_forward][row][column]
                difference_here = threshold - value_here
                percentage = difference_here/difference *100 
                reduction_matrix[row][column] = 255 - value_here
                
            else:
                reduction_matrix[row][column] = -1 #-1 is placeholder. 
    return reduction_matrix


# In[165]:


#Use np.shape() to get 0 = slice, 1 = rows, 2 = columns.
def extract_all_TTC_values(Stack, TimeArray, time_increment, start_time):
    shape = np.shape(Stack)
    data = pd.DataFrame() #empty dataframe
    time = np.zeros(shape[0])
    new_pixels = np.zeros(shape[0]) ##Should be an empty array the size of shape[0] ie all slices. Want to store
    #the running_number_of_pixels count for each timepoint here. 

    #//make a time column 
    time = [] #empty list, which needs to be populated. 
    for timepoint in range(shape[0]):
        time_here = start_time + time_increment*timepoint
        time.append(time_here)

    #made a time column, add it to the dataframe. 
    data["Time (hr)"] = time
    
    for images in range(shape[0]): 
        running_TTC_value = np.zeros(len(Stack[images:])) #np.zeros makes an array of zeros for every slice 50 and later. (number of slices - starting index)
        running_number_of_pixels = 0 #this is just an integer.
        for rows in range(shape[1]): #loop thorugh all rows. 
            for columns in range(shape[2]): #loop through all of the columns
                if TimeArray[rows][columns] == images:
                    TTC_value_here = Stack[images:, rows, columns] #an array of TTC values for one pixel where each subsequent value is the TTC value in the next slice. 
                    running_TTC_value = TTC_value_here + running_TTC_value
                    running_number_of_pixels = running_number_of_pixels + 1 #(or running_number_of_pixels +=1)
        new_pixels[images] = running_number_of_pixels
                    
                
        #// calculating average TTC value in every loop of time. 
        if running_number_of_pixels > 0:
            average_TTC_value = running_TTC_value/running_number_of_pixels
        else:
            average_TTC_value = running_TTC_value #Exact same as storage.
            
        #// need to store it and keep it the same length in every loop. 
        #this is where we need to pad with zeros of negative ones - needs to be the same size as time. 
        storage = np.full(images, -1) #makes an array of -1's with a negative 1 for every slice. 
        #want to take the average_TTC_values and put it into storage, overwriting at the correct starting position. 
        storage= np.append(storage, average_TTC_value)  
        data[f"Pixels which appear at slice number {images} (image)"] = storage
    return data, new_pixels        


# In[166]:


def extract_all_TTC_values_SD(Stack, TimeArray, time_increment, start_time):
    shape = np.shape(Stack)
    data = pd.DataFrame() 
    time = []
    new_pixels = np.zeros(shape[0]) 
    # Converting slice numbers into hour values.
    for timepoint in range(shape[0]):
        time_here = start_time + time_increment*timepoint
        time.append(time_here)
    # Adding time values (in hours) to DataFrame.
    data["Time (hr)"] = time
    
    # New additions to the function from here:
        
    # 'images' in this for loop refers to each slice in the stack.
    # TTC and new pixel values are for each slice, not cumulative.
    for images in range(shape[0]): 
        running_number_of_pixels = 0 
        # List of lists to store all TTC values over time for all new pixels.
        all_TTC_values = []
        for rows in range(shape[1]): 
            for columns in range(shape[2]):
                if TimeArray[rows][columns] == images:
                    # TTC values over time for that pixel are appended to the
                    # list of lists for all new pixels (all_TTC_values).
                    all_TTC_values.append(list(Stack[images:, rows, columns]))
                    running_number_of_pixels += 1
        
        # Saving the number of new pixels for the function's first output.
        new_pixels[images] = running_number_of_pixels
        
        # Initialising lists for TTC average and SD values at each timepoint.
        average_TTC_list = []
        TTC_SD_list = []
        
        # Iterating through all timepoints after the 'images' slice.
        for timepoint in range(shape[0]-images):
            # New list containing all TTC values at that timepoint (from each
            # pixel's TTC value list in the 'all_TTC_values' list of lists).
            TTC_at_timepoint = [pixel_TTC_list[timepoint] for pixel_TTC_list 
                                in all_TTC_values] 
            # Calculating/appending average TTC and SD at that timepoint.
            average_TTC = np.mean(TTC_at_timepoint)
            average_TTC_list.append(average_TTC)
            TTC_SD = np.std(TTC_at_timepoint)
            TTC_SD_list.append(TTC_SD)
        
        # Front-loading the average TTC and SD lists with 'nan' so that they
        # start at the correct timepoint within the DataFrame.
        while len(average_TTC_list) != shape[0]:
            average_TTC_list.insert(0, np.nan)
        while len(TTC_SD_list) != shape[0]:
            TTC_SD_list.insert(0, np.nan)
        
        # Adding the average TTC and SD values to the DataFrame.
        average_TTC = pd.Series(average_TTC_list)
        TTC_SD = pd.Series(TTC_SD_list)
        data[f"Average TTC: Appearing at slice {images}"] = average_TTC
        data[f"SD: Appearing at slice {images}"] = TTC_SD
        
    return data, new_pixels 


# In[167]:


#First load the images of the movie and generate a stack for the script to read.
    #ex. movie_stack = generate_stack("folder_name_containing_images")
#In our case, we filter the images so we are analyzing the images for every hour, instead of every 30 minutes.
    # ex. movie_stack_filter = filter_movie(movie_stack,2)
#Then we generate a time matrix which stores the values of when the pixels entered the colony. 
    #ex. movie_stack_time_matrix = generate_time_matrix(movie_stack_filter, 160) 
#The refined threshold will be printed for manual confirmation. 


# In[168]:


# Plotting the time matrix is helpful in determining if there were any issues. 
    #ex. plt.imshow(movie_stack_time_matrix)


# In[169]:


# Then we extract the average pixel value for all of the pixels which were detected at a given timepoint. 
    #ex. movie_stack_TTC, new_pixels = extract_all_TTC_values_SD(movie_stack_filter, movie_stack_time_matrix, 1, 0)


# In[170]:


#Then save the file(s) as a csv. 
    #ex. moive_stack_TTC.to_csv("desired_file_name.csv")
